# PPARG in osteocytes controls cell bioenergetics and systemic energy metabolism independently of sclerostin levels in circulation

**DOI:** 10.1101/2024.04.04.588029

**Authors:** Sudipta Baroi, Piotr J. Czernik, Mohd Parvez Khan, Joshua Letson, Emily Crowe, Amit Chougule, Patrick R. Griffin, Clifford J. Rosen, Beata Lecka-Czernik

## Abstract

**Objective:** The skeleton is one of the largest organs in the body, wherein metabolism is integrated with systemic energy metabolism. However, the bioenergetic programming of osteocytes, the most abundant bone cells coordinating bone metabolism, is not well defined. Here, using a mouse model with partial penetration of an osteocyte-specific PPARG deletion, we demonstrate that PPARG controls osteocyte bioenergetics and their contribution to systemic energy metabolism independently of circulating sclerostin levels.

**Methods:** *In vivo* and *in vitro* models of osteocyte-specific PPARG deletion, i.e. *Dmp*1^Cre^*Pparγ*^flfl^ male and female mice (γOT^KO^) and MLO-Y4 osteocyte-like cells with either siRNA-silenced or CRISPR/Cas9-edited *Pparγ*. As applicable, the models were analyzed for levels of energy metabolism, glucose metabolism, and metabolic profile of extramedullary adipose tissue, as well as the osteocyte transcriptome, mitochondrial function, bioenergetics, insulin signaling, and oxidative stress.

**Results:** Circulating sclerostin levels of γOT^KO^ male and female mice were not different from control mice. Male γOT^KO^ mice exhibited a high energy phenotype characterized by increased respiration, heat production, locomotion and food intake. This high energy phenotype in males did not correlate with “beiging” of peripheral adipose depots. However, both sexes showed a trend for reduced fat mass and apparent insulin resistance without changes in glucose tolerance, which correlated with decreased osteocytic responsiveness to insulin measured by AKT activation. The transcriptome of osteocytes isolated from γOT^KO^ males suggested profound changes in cellular metabolism, fuel transport and usage, mitochondria dysfunction, insulin signaling and increased oxidative stress. In MLO-Y4 osteocytes, PPARG deficiency correlated with highly active mitochondria, increased ATP production, shifts in fuel utilization, and accumulation of reactive oxygen species (ROS).

**Conclusions:** PPARG in male osteocytes acts as a molecular break on mitochondrial function, and protection against oxidative stress and ROS accumulation. It also regulates osteocyte insulin signaling and fuel usage to produce energy. These data provide insight into the connection between osteocyte bioenergetics and their sex-specific contribution to the balance of systemic energy metabolism. These findings support the concept that the skeleton controls systemic energy expenditure *via* osteocyte metabolism.

**Highlights:**

- Osteocytes function as a body energostat via their bioenergetics
- PPARG protein acts as a “molecular break” of osteocyte mitochondrial activity
- PPARG deficiency activates TCA cycle, oxidative stress and ROS accumulation
- PPARG controls osteocyte insulin signaling and fuel utilization

## 1. Introduction

The nuclear receptor and transcription factor PPARG is a known regulator of metabolic multiverse on the systemic and cellular level including bone metabolism reflected by the dynamics of bone remodeling. The research on the skeletal function of PPARG started in the late 1990s and was fueled by the discovery of the inverse relationship between osteoblast and adipocyte differentiation driven by this transcription factor, and its role in the regulation of osteoclast differentiation [1–6]. Previously, we demonstrated that PPARG is highly expressed in osteocytes, where it directly controls *Sost* gene expression *via* multiple PPREs present in the proximal and distal promoter region, and PPARG natural and pharmacologically induced activities positively correlate with the levels of sclerostin protein in bone [7,8]. Recently, it has been demonstrated that osteocytic PPARG controls metabolic function of extramedullary adipose tissues *via* sclerostin, which as an inhibitor of WNT pathway and augments adipocyte development with lipid storing function [9]. The same study showed that low levels of sclerostin in circulation induce beiging of extramedullary white adipose tissues (WAT). Another study identified osteocytic PPARG as a positive regulator of BMP7 production, a cytokine which can act as an endocrine circulating factor to increase extramedullary fat metabolism [10]. Together, both studies are consistent in demonstrating that PPARG deletion from cells of osteoblast/osteocyte lineage leads to a greater energy metabolism phenotype associated with increased glucose metabolism and insulin sensitivity, as well as beiging of epididymal and inguinal WAT [9,10].

The skeleton contributes to metabolic homeostasis through the process of bone remodeling which is fundamental for maintenance of bone mass and quality, but requires significant energy. Bone remodeling consists of synchronized steps of bone resorption by osteoclasts and subsequent bone formation by osteoblasts. This is a metabolically demanding process that influences systemic energy balance, as it entails a continuous supply of energy metabolites to the bone in the form of glucose and fatty acids. Glucose is a major energy source used for differentiation and function of osteoblasts and osteoclasts. In these cells, ATP production from glucose can be complemented with oxidative metabolism of fatty acids, and to a much lower extent glutamine, and the source of fuel aligns with osteoblast differentiation, maturation and bone forming activity [11–14]. Whereas osteoblast and osteoclast energy needs are relatively defined, the metabolic status of osteocytes, and how that relates to their function, is in the early stages of elucidation, in part due to relative limitations in applying established experimental techniques to osteocytes in their *in vivo* biologic environment (reviewed in [15]).

Osteocytes are considered the major endocrine cells in bone. They constitute 90-95% of bone cells and are localized in the mineralized bone compartment. It is estimated that in humans the number of osteocytes comprises up to 40 billion cells, roughly half of the number of brain neurons, with total length of dendritic processes matching that in the brain [16]. Therefore, even subtle changes in osteocyte metabolism or production of secreted proteins may have significant local and systemic effects. Osteocytes orchestrate bone remodeling by producing factors regulating bone formation by osteoblasts and bone resorption by osteoclasts. Among the best characterized, osteocytes produce sclerostin which inhibits bone formation by inhibiting the WNT pathway activity in osteoblasts, and RANKL which is indispensable for osteoclast differentiation and bone resorption [17].

The Cre-LoxP system is commonly used to create animal models with genetic alteration in expression of specific genes and proteins in selected type of cells. This advance in genetic manipulation of animal models has been incredibly useful in uncovering new physiologic and pathologic processes and development of therapeutic means targeting them with relative precision. However, the Cre-LoxP system is not ideal and suffers from unpredictable flaws which may inadvertently affect primary and secondary cellular and whole-body mechanisms. The prevailing issue lies within the Cre-LoxP germline models, in which upon chromosomal insertion Cre recombinase may lose activity to some extent and over time [18]. To get around that concern and to focus on other metabolic factors that may arise from osteocytes, we took advantage of our mouse model with a partial deletion in osteocytic PPARγ (γOT^KO^) in which circulating sclerostin was not affected. This partial model unveiled unexpected contribution of osteocytes to the systemic energy metabolism which had not been detected in other similar models with more efficient PPARG deletion from osteocytes. In this report we demonstrate that mitochondrial dysfunction of PPARG-deficient osteocytes leads to increased oxidative stress and accumulation of ROS, which could potentially have long-term implications for skeletal health. We conclude that osteocyte bioenergetics is an essential component of systemic energy balance and, at least in males, osteocytes under PPARG control act as systemic energostat in this process.

## 2. Materials and Methods

### 2.1 Animals

Osteocyte-specific PPARG knock-out mice (*Dmp*1^Cre^*Pparγ*^flfl^ or γOT^KO^) were developed at the University of Toledo, by crossing Dmp1-Cre (Stock No: 023047, The Jackson Laboratory, Bar Harbor, ME) with Pparγ^loxP^ (Stock No: 004584, The Jackson Laboratory) as previously described [7]. In all experiments, littermates with *Dmp*1^Cre^*Pparγ*^+/+^ genotype were used as control (Ctrl). The animals were maintained under 12hrs dark-light cycle with ad libitum access to water and chow, either regular (Teklad global 16% protein rodent diet; code:2916) or breeding (Teklad global 19% protein extruded rodent diet: code:2919). The breeding and experimental protocols (#107229 and #105923, respectively) conformed to the NIH National Research Council’s Guide for the Care and Use of Laboratory Animals and were reviewed and approved by the University of Toledo Health Science Campus Institutional Animal Care and Utilization Committee. The University of Toledo animal facility is operating as a pathogen-free, AAALAC approved facility and animal care and husbandry meet the requirements in the Guide for the Care and Use of Laboratory Animals.

### 2.2 Measurements of body composition and indirect calorimetry

Measurements of body composition and indirect calorimetry of experimental animals started at 2 mo of age and continued on the same groups of mice for up to 6 mo of age. The minispec mq7.5 NMR analyzer (Bruker) and Bruker minspec software v2.58 were used for the measurements and calculations. Indirect calorimetry was performed using Oxymax Comprehensive Lab Animal Monitoring System (CLAMS) (Columbus Instruments, Columbus OH) with cage-set-up for 16 animals. For both males and females, the experimental groups consisted of 5 γOT^KO^ and 11 Ctrl mice. The measurements included food and water intake, oxygen consumption, carbon dioxide release, respiratory exchange ratio (RER), heat generation, and physical activity (planer, planer ambulatory and vertical). After one day of cage acclimatization, the measurements were performed for the next three consecutive days. Measurement values were normalized by animal lean mass assessed by NMR, as described above.

### 2.3 Intraperitoneal Glucose Tolerance Test (GTT) and Insulin Tolerance Test (ITT)

GTT and ITT were performed on γOT^KO^ (n=5) and Ctrl mice (n=11) at 4- and 6-months of age. Mice were fasted for 4 hr before assays. For GTT, they were injected peritoneally with sterile filtered 20% isotonic glucose solution at a dose of 2 g/kg body weight, while for ITT they were injected with insulin (Humulin; Healthwarehouse, Cat#A10083401) at a dose of 0.75 units/kg body weight. Glucose levels were measured in blood after tail nipping at indicated time intervals including time 0 recorded immediately before injection. Measurements were done using Sunmark TrueTrack glucometer (Nipro Diagnostics. Cat# 797461)) and Sunmark TrueTrack blood glucose test strips (Trividia Health, Cat#56151-0810-01).

### 2.4 Primary cells and cell lines

Isolation of primary osteocytes is described in [7]. RNAs and proteins were isolated immediately after osteocyte liberation from bone matrix and used for transcriptome analysis, as described below. Osteocyte-like MLO-Y4 cells were cultured on collagen coated plates at 37°C with 5% CO2 supply in the presence of alpha-MEM media supplemented with 5% fetal bovine serum (FBS), 5% calf serum (CS), and 1% penicillium/streptomycin (P/S).

### 2.5 siRNA silencing (γY4^KD^) and CRISPR/Cas9 editing of Pparγ (γY4^KO^) in MLO-Y4 cells

PPARG silencing was achieved by using *Pparγ* siRNA (Santa Cruz Biotechnology; Cat#sc43530) with DsiRNA (Integrative DNA Technologies, Coralville, IA; Cat# 51-01-14-03) used as a negative control. RNA oligonucleotides were delivered to MLO-Y4 cells using the X-treme Gene siRNA Transfection Reagent (Roche; Cat#04476093001) according to a protocol provided by the manufacturer. Cells were used for assays 36 hrs after transfection.

Alternatively, PPARG was knocked-out in MLO-Y4 cells with CRISPR/Cas9 editing system designed by Synthego CRISPR Gene Knockout Kit v2 (Synthego Corporation, Redwood City, CA; Cat # SO6765399). Three guide RNAs, SEQ1: G*A*G*AAAUCAACUGUGGUAAA, SEQ2: A*G*A*GCUGAUUCCGAAGUUGG, SEQ3: U*U*C*CACUUCAGAAAUUACCA, were used to make ribonucleoprotein (RNP) complex with Cas9 2NLS (nuclear localization signal) in 1.3:1 ratio. The stars indicate 2’-O-methyl analogs and 3’-phosphorothioate internucleotide linkages. Transfection was performed using Lipofectamine CRISPRMAX transfection reagent (Invitrogen, Cat#CMAX00003). Briefly, MLO-Y4 cells were trypsinized, counted and volume adjusted to 60,000 cells in 100 µl OptiMEM. For each well of a 24-well plate, 50 µl of transfection solution (RNP complex + transfection reagent) was added to 100 µl of cell suspension, mixed gently and allowed to stand for 10 min before plating to the wells containing 150 µl OptiMEM warmed up to 37C. Four hours post transfection, 300 µl Alpha-MEM containing 10% FBS, 10% CS, and 1% P/S was added to each well. As a negative control, transfection was performed without addition of 2NLS. As a positive control, transfection was performed using Rosa26 guide RNA (5’-GAGGCGGATCACAAGCAATA-3’). Seventy-two hours post-transfection, DNA was isolated using Trizol for analysis of *Pparg* gene sequence editing.

Specific primers were designed covering 442 bp region around the edited site (F: TCAGGAAACCAGATGCCACA, R: TCAGCCTAAGACAAAACTGGCA) were used for PCR amplification using Taq polymerase. Amplicons were purified using Qiagen PCR purification kit (Cat#28104). 1% Agarose gel was run with amplicons to confirm a single band. Purified PCR amplicons were then sequenced (Sequencing primer 5’-GTGTGTGTGTGTAAGTTTGGGAAC-3’, 25 pmol for 10 ng purified PCR product) by company Genewiz, NJ, USA. The sequences were analyzed using ICE (Interference of CRISPR Edits), an online bioinformatic software provided by Synthego to evaluate editing (KO) efficiency.

The transfected cell samples with highest KO score were used for clonal selection. After trypsinization, cells were plated on collagen coated 96-well plates in the dilution of single cell per well (7 plates). Cultures were monitored for colony formation from 2 to 8 weeks after plating using Incucyte live cell imaging system (Sartorius). Wells with single colonies were harvested and cell cultures were expanded for making stocks and for DNA isolation, first on 12 well plates followed by 6 well plates. Isolated DNA were processed as mentioned before with PCR and sequencing and the KO efficiency was analyzed using ICE.

### 2.6 Transcriptomic analysis of primary osteocytes and MLO-Y4 CRISPR Pparγ KO (γY4^KO^) clones

Osteocyte RNA from γOT^KO^ (n=4) and Ctrl (n=3) 4 mo old male mice was isolated using TRIZOL and transcriptomic analysis was performed by Arraystar Inc. (Rockville, MD, USA) using the Mouse LncRNA Array v4.0 platform (8 x 60K, Arraystar, Inc). To increase stringency, the same RNA was also analyzed with Next Generation Sequencing (NGS) using flow of NovaSeq SP 100 cycles (Wayne State University Bioinformatics Core, Detroit, MI). Similarly, transcriptomes of γY4^KO^ clones with highest KO scores were analyzed by NGS, as above. Raw data files are deposited in the NCBI Gene Expression Omnibus repository (GEO), submission number: (pending).

From the curated data, transcripts with more than a 2-fold change and a p-value less than 0.05 were recognized as differentially expressed, and the GO term enrichment and network enrichment analysis were performed. To increase the degree of confidence on the bioinformatic analysis of GO term enrichment and for comparison, a parallel analysis was performed using g:GOSt program of opensource enrichment analysis web server g:Profiler [19]. Along with BP (Biological Process), MF (Molecular Function) and CC (Cellular Component), g:Profiler allows for network enrichment using Reactome and KEGG database. In addition, g:Profiler provided TF (Transcription Factor) enrichment analysis.

### 2.7 Seahorse Mito-Stress test and Mito-Fuel Flex test

For Mito-Stress test, MLO-Y4 cells with *Pparγ* silenced with siRNA (γY4^KD^) and their scrambled control (Scrl) were plated 6 hrs before the assay onto Seahorse cell culture microplates (Agilent Technologies, Santa Clara, CA; Cat#101085-004). Four corner wells of the plate were used as negative control. Assay medium consisted of Sea Horse XF base medium without phenol red (Agilent Technologies, Cat#103335-100) supplemented with glucose (25 mM), sodium pyruvate (1 mM) and glutamine (2 mM). Mitochondria function was tested in the presence of stressors such as Oligomycin (1 µM), FCCP (2 µM), and Rotenone/Antimycin (0.9 µM) to block ATP-synthase, uncoupling, and shutting down entire oxidative phosphorylation process, respectively. Oxygen consumption rate (OCR) and extracellular acidification rate (ECAR) were measured at basal level and after addition of stressors using Seahorse XFe96 analyzer (Agilent Technologies). Assay operation software Wave version 2.6.1 was used for running the assay and obtaining data. For normalization, live cells stained with Hoechst 33342 (1:2,000 dilution) (ThermoScientific, Cat#62249) were enumerated using Cytation 5 plate reader (BioTek Instruments - Agilent). Values obtained from the Mito-Stress test were normalized per 1,000 cells.

Fuel preference of CRISPR/Cas9 *Pparγ* edited cells (γY4^KO^) and mock-edited cells (C) was measured using Seahorse Mito-Fuel Flex test (Agilent Technologies), in assay medium as above and according to the manufacturer protocol. Cells were plated on assay micro-plate at the density of 20,000 cells per well. The assay uses three inhibitors: UK5099 (an inhibitor of the glucose oxidation pathway), BPTES (an inhibitor of the glutamine oxidation pathway), Etomoxir (an inhibitor of long chain fatty acid oxidation) provided in the Seahorse XF Mito fuel flex test kit (Agilent Technologies, Cat#103260-100). Fuel dependency for a specific pathway was measured after inhibiting this pathway. Fuel capacity of that pathway was measured after inhibiting two other pathways with specific inhibitors. Each measurement was performed on 8 replicate wells. Software Wave version 2.6.1 was used for running the assay and obtaining data.

### 2.8 TMRE-based and ERthermAC-based measurements of mitochondrial activity

Mitochondrial membrane potential was measured in γY4^KD^ cells with *Pparγ* silenced with siRNA and their scrambled control (Scrl) using Mitochondrial Membrane Potential Assay kit (Abcam, Cat# ab113852). Cells were plated on collagen coated 24-well plate at density of 50,000 cells/well (5 well/per group). Twelve hours after plating, cells were stained with TMRE following protocol provided by manufacturer. Fluorescent intensity of TMRE labelled cells were measured using Cytation 5 plate reader at 549/575 nm excitation/emission wavelength. Data were normalized after correction for background originating from intrinsic fluorescence of collagen.

Optical visualization of thermogenesis was performed using ERthermAC fluorescence dye in accordance to the published protocol (20). Cells were grown on collagen-coated 24-well plates to approximately 80% confluence in αMEM medium supplemented with 5% FBS, 5% CS and 1% P/S. Cells were stained in separate wells using fresh medium aliquots containing ERthermAC dye at 250 nM, and Rhodamine B at 6.3 μM final concentration for 20 minutes at 37^ο^C (MilliporeSigma). Images of cells were acquired with Incucyte S3 Live-Cell Analysis System at 15 minutes intervals using 20X magnification in red channel (567-607 nm excitation, 622-704 emission). Images recorded at 0.625 μm resolution were analyzed using ImageJ software. Image pre-processing included background subtraction using rolling ball algorithm at 100 μm diameter, and thresholding of color pixel brightness empirically set to 15-255 value range for all analyzed images. Object size was set to a range of 100-10000 pixels that was the most representative object size observed across all analyzed images. Mean pixel density was used as a signature of temperature-dependent change in fluorescence emitted by a single cell.

### 2.9 Gene expression analysis using quantitative real-time RT-PCR

Total RNA from different fat depot was isolated using TRIzol (Thermo Fisher Scientific, Cat# 15596026). One-half μg of RNA was converted to cDNA using the Verso cDNA synthesis kit (Thermo Fisher Scientific, Waltham, MA). PCR amplification of the cDNA was performed by quantitative real-time PCR using TrueAmp SYBR Green qPCR SuperMix (Smart Bioscience, Maumee, OH) and processed with StepOne Plus System (Applied Biosystems, Carlsbad, CA). The thermocycling protocol consisted of 10 min at 95°C, 40 cycles of 15 s at 95°C, 30 s at 60°C, and 20 s at 72°C, followed by melting curve step with temperature ranging from 60 to 95°C to ensure product specificity. Relative gene expression was measured by the comparative CT method using 18S RNA levels for normalization. Primers were designed using Primer-BLAST (NCBI, Bethesda, MD) and are listed in Supplementary Table 1.

### 2.10 Diagnostic assays

To measure response to insulin, the CRISPR/Cas9-edited γY4^Ctrl^ and γY4^KO^ were grown in αMEM media supplemented with 10% FBS and 1% P/S until cultured achieved 80% confluency. Media were replaced with serum-free αMEM supplemented with 1% P/S and 20 mM HEPES for 2 hrs followed by media change to the same media with or without 100 nM insulin. After 30 min of incubation, cells were harvested for protein lysate using lysis buffer (20nM Tris pH 7.5, 150mM NaCl, 1mM EDTA, 1% Triton (by weight), 2.5mM sodium pyrophosphate, 1mM beta-glycerophosphate, supplemented with protease and phosphatase inhibitors). Protein concentration was measured using the BCA assay (Thermo Fisher Scientific, Waltham, MA 23225). Fifteen µg of lysate in Laemmli sample buffer was heated at 95°C for 5 minutes and loaded onto a 10% SDS-PAGE gel and ran at 115 volts for 1-1.5 hours. Proteins were transferred to a PVDF membrane using a tank transfer system at 100 volts for 1 hour. The membrane was blocked with 5% BSA for 1 hour and washed 3 times for 5 minutes in 1X TBST. Primary antibody incubation occurred overnight at 4°C (AKT, p-AKT, PPARG, and β-Actin dilutions were all 1:1,000 in 5% BSA). Membranes were washed as before and incubated in secondary antibody at a 1:10,000 dilution for 1 hr. Membranes were washed and developed using the ECL method (Thermo Fisher Scientific, Waltham, MA 34577). Imaging was performed on a Syngene GBOX Chemi XX6 (Syngene, Frederick, MD 21704). Antibodies used for Western blots: PPARG Rb monoclonal (cat#81B8; Cell Signaling Technologies), AKT Rb polyclonal (cat#9272S; Cell Signaling Technologies), Phospho-AKT (Ser 473) Rb polyclonal (cat#9271S; Cell Signaling Technologies), β-Actin Ms monoclonal (A1978; Sigma), Anti-Rabbit IgG HRP-linked (cat#7074; Cell Signaling Technologies), and Anti-Mouse IgG HRP-linked (cat#sc-2005; Santa-Cruz Biotechnology).

Adiponectin levels in serum of γOT^KO^ and Ctrl mice were measured using ELISA (Quantikine ELISA, R & D systems, Cat#MRP300). Each sample measurement was performed in duplicate using 10 µl serum. Serum insulin, cholesterol, and triglyceride levels in γOT^KO^ and Ctrl mice were analyzed by Chemistry Core of Michigan Diabetes Research Center (University of Michigan Ann Arbor, MI). To measure oxidative stress and reactive oxygen species (ROS) cellular activities two assays were used. The GSH/GSSG Ratio Detection Assay (Abcam cat# ab138881) measured conversion of glutathione from reduced stage (GSH) to the oxidized stage (GSSG) upon exposure to oxidative stress. An increased ration of GSSG-to-GSH is an indicator of oxidative stress. The ROS activities were quantitated with DCFDA/H2DCFDA – Cellular Reactive Oxygen Species Detection Assay (Abcam cat# ab113851). Both assays were used according to protocol recommended by manufacturer.

### 2.12 Statistical analysis

Data are represented as means ± SD and were analyzed by statistical analysis software GraphPad Prism v.9. Students t-test was used to compare statistical significance between two groups and ANOVA was used to compare between more than two groups. p-value less than 0.05 was considered as statistically significant.

## 3. Results

### 3.1 γOT^KO^ male mice have increased systemic energy metabolism regardless of sclerostin levels in circulation

Recent findings have demonstrated a critical role of osteocytic sclerostin in regulation of metabolism of extramedullary fat depots [9]. Thus, decreased sclerostin levels in circulation lead to peripheral white adipose tissue (WAT) beiging which in consequence changes levels of systemic energy metabolism. We reported previously that PPARG acts as a positive transcriptional regulator of sclerostin expression in osteocytes and its deletion specifically in osteocytes correlates perfectly with a lack of sclerostin production (Pearson Correlation R^2^= 0.982) [7]. In that study and the study presented here, we used a murine model of osteocyte-specific deletion of PPARG protein, resulting from crossing 10 kb Dmp1^Cre^ and Pparγ^loxP^ mice (γOT^KO^) to delete exon 1 and 2 from *Pparγ* gene sequence. However, as presented previously and shown here, in this model we have not seen significant changes in the levels of circulating sclerostin in both males and females, probably due to not complete penetration of PPARG KO phenotype, as it is discussed later (Fig. 1A) [7].

**Figure 1.**
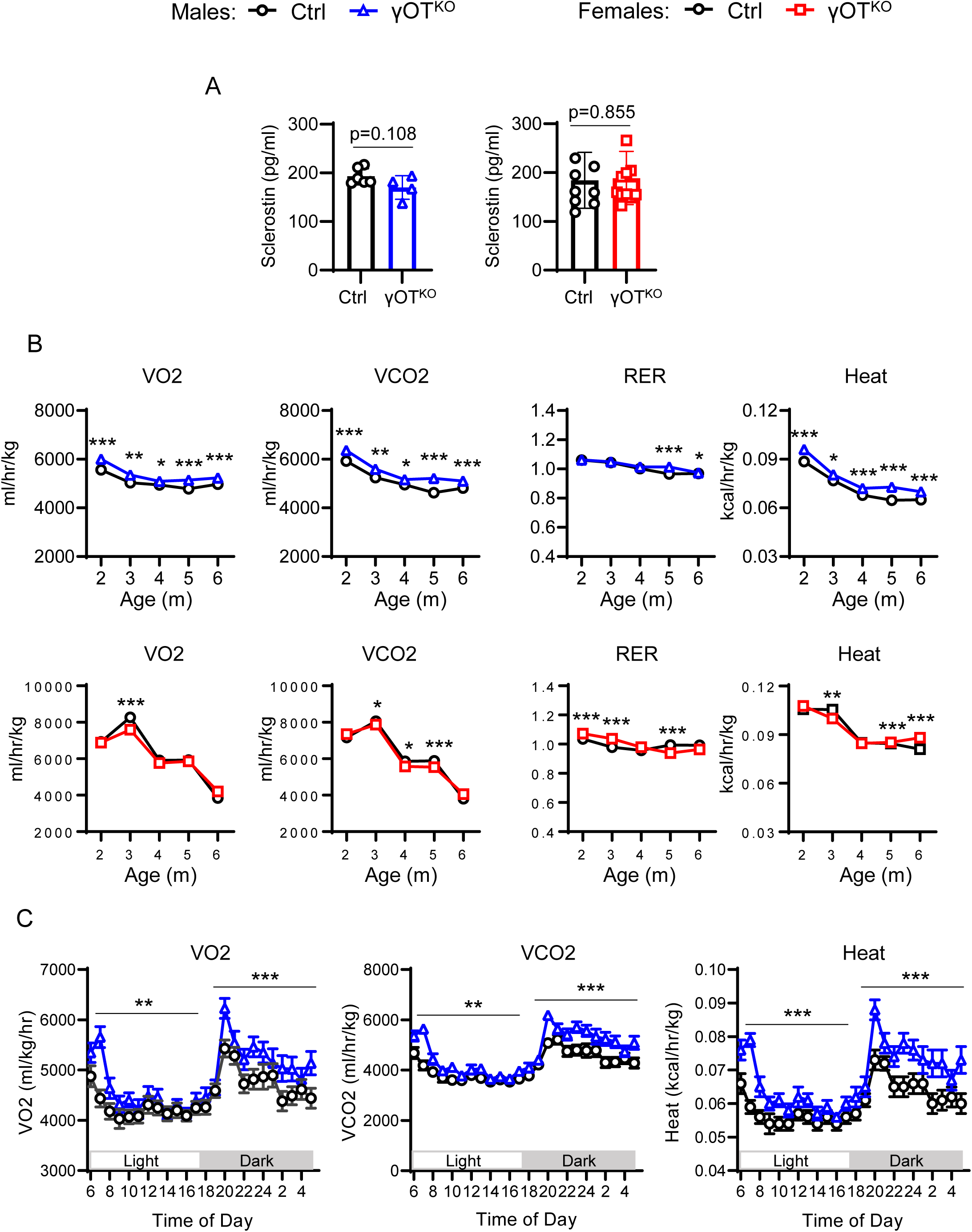
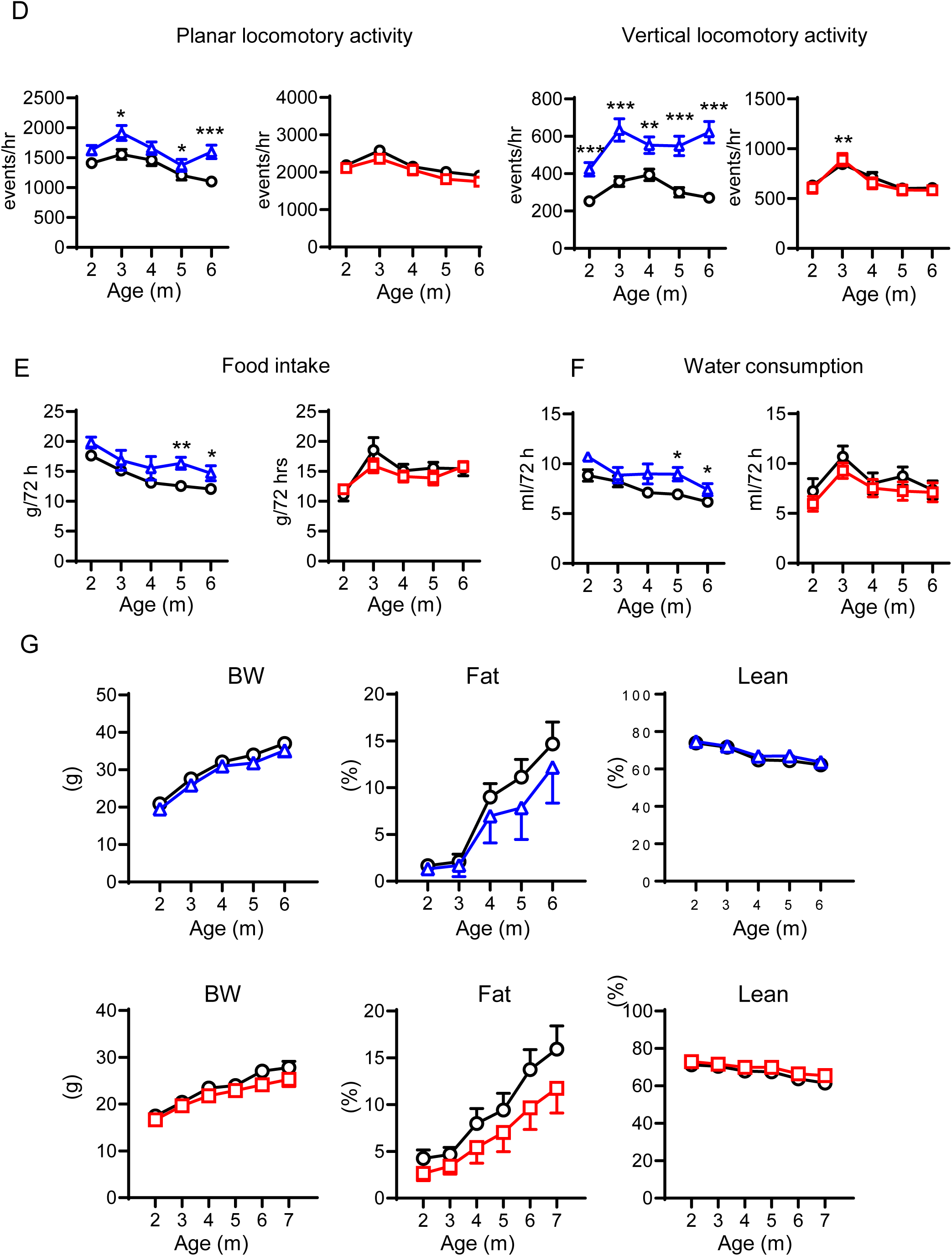
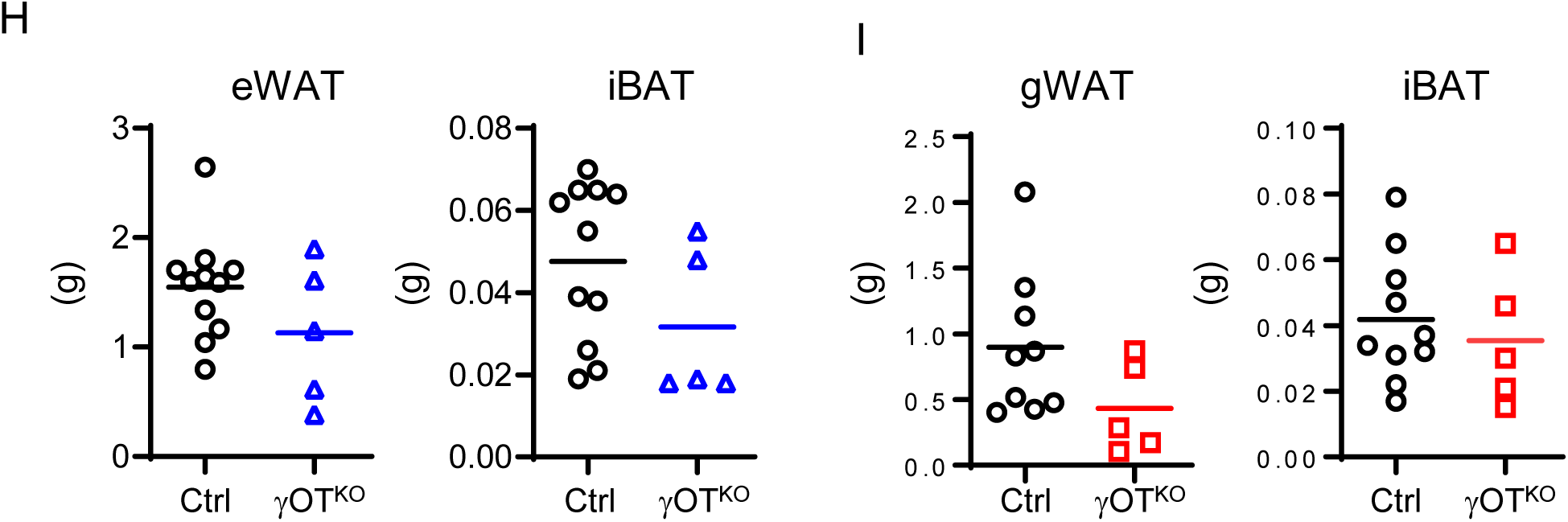
Sclerostin levels in circulation and metabolic parameters of γOT^KO^ male and female mice. A. Sclerostin levels measured in serum of male and female mice at age 6 – 10 mo old (males – Ctrl: n=6, γOT^KO^=4; females – Ctrl: n=8, γOT^KO^=11). B. Longitudinal measurements of respiration in light (12 hr) and dark (12 hr) day cycles using the Comprehensive Lab Animal Monitoring System (CLAMS). Each point represents an average of 3 days measurements in males (blue) and females (red). VO2 - oxygen consumption, VCO2 – carbon dioxide production, RER - Respiratory Exchange Ratio. C. Hourly respiration and heat production of 6 mo old males monitored for 24 hrs during light and dark day cycle. D. Average locomotory activity measured during 3 days of CLAMS measurements (n=4 animals per each group). E and F. Food and water consumption during 72 hrs of CLAMS measurements at each indicated age. G. Body weights (BW) and body composition measured by NMR at the time of CLAMS experiments. H. Weights of epidydimal WAT (eWAT) and interscapular BAT (iBAT) of 6 mo old males. I. Weights of gonadal WAT (gWAT) and interscapular BAT (iBAT) of 6 mo old females. If not differently specified all measurements included: males - age: 2m – 6m, Ctrl: n=11, γOT^KO^: n=5 mice; females - age: 2m – 6m, Ctrl: n=11, γOT^KO^: n=5 mice. For A, E – I, unpaired two-tailed student’s t-test was used for statistical comparison of two experimental points and groups. For B and D statistical significance of differences between groups were calculated using non-parametric Mann-Whitney test. For C statistical significance was calculated using paired student’s t-test comparing each time points of light and dark cycles. p < 0.05 was considered significant. *p < 0.05; **p < 0.01; ***p < 0.001.

The metabolic phenotype of γOT^KO^ mice had been assessed as a function of sex and age. The same cohort of male and female mice had been monitored on a monthly basis for body and metabolic parameters from the age of 2 to 6 month. In contrast to littermate γOT^KO^ female mice, male γOT^KO^ mice consistently demonstrated increased respiration and increased energy expenditure. Indirect calorimetry using CLAMS metabolic cages system, showed increased oxygen consumption (VO2) and increased carbon dioxide production (VCO2), especially during day dark cycle when animals are naturally more active (Fig. 1B and 1C). Higher VO2 and VCO2 were observed as early as 2 mo of age and persisted throughout the length of the study up to 6 mo of age. In contrast, γOT^KO^ female mice showed modest but significant decrease in respiration (Fig. 1B). RER was not much different in younger γOT^KO^ males and slightly increased with aging suggesting increase in carbohydrate metabolism. In contrast, higher RER was observed in younger γOT^KO^ females and become slightly lower with age suggesting an increase in lipid metabolism (Fig. 1B).

The γOT^KO^ animals generated more heat. In males, an increase in heat production started at 2 mo of age and continued throughout observation period, while in females increased heat production was delayed and become higher in 5 and 6 mo old animals (Fig. 1B). The most stunning difference between sexes was observed in the animals actigraphy. The γOT^KO^ males showed high planar (X-plane associated with eating and rummaging) and vertical (Z-plane associated with drinking and surveying) activities at 2 mo of age which continued with advancing age, while females locomotory activities were not changed (Fig. 1D). These data suggest the role of osteocytic PPARG in regulation of systemic energy metabolism and sexual divergence in this regulation.

### 3.2 Body composition and peripheral fat function are mostly unaffected in γOT^KO^ mice

The respiratory and actigraphy profile showed in Fig. 1 corresponded to higher food consumption and water intake in γOT^KO^ males, and not different in females (Fig. 1E and 1F). Body weight of γOT^KO^ males and females were not significantly different from their respective Ctrl at any analyzed time points (Fig. 1G). However, in both sexes the tendency to lower fat contribution to the overall body composition had been observed. There was no significant difference in the weight of epididymal or perigonadal white adipose tissue (eWAT and gWAT, respectively) and interscapular brown adipose tissue (iBAT) isolated from 6 mo old male and female γOT^KO^ mice, although a tendency of decreased weight of fat depots was observed, as compared to littermate Ctrl mice (Fig. 1H and 1I).

Gene expression analysis of three different adipose tissue depots which contribute significantly to the levels of systemic energy metabolism, namely eWAT/gWAT, iBAT, and inguinal WAT (iWAT) which is known for its capabilities to convert to beige fat upon hormonal or pharmacological treatment, did not show fat beiging in γOT^KO^ mice, as it was demonstrated previously in similar models [9,10]. In general, the expression of beige phenotype markers: *Ucp1*, *Prdm16*, and *Dio2*, was not different in eWAT/gWAT and iBAT in males and females as compared to Ctrl, with minor exceptions (Fig. 2A and 2B). *Ucp1* expression, but not other markers, was elevated in gWAT of 6 mo old γOT^KO^ females, while *Dio2* expression was elevated in iBAT of 6 mo old γOT^KO^ males. The expression of beige markers was not affected in males and females iWAT (Fig. 2C). The expression of two adipokines essential for maintenance of energy metabolism balance, adiponectin and leptin, was not affected in eWAT/gWAT and iBAT of γOT^KO^ males and females but was decreased in iWAT of γOT^KO^ males as compared to Ctrl (Fig. 2C). These indicated no changes in the metabolic activities of adipocytes in these fat depots and possible decrease in endocrine activities of iWAT in males.

**Figure 2.**
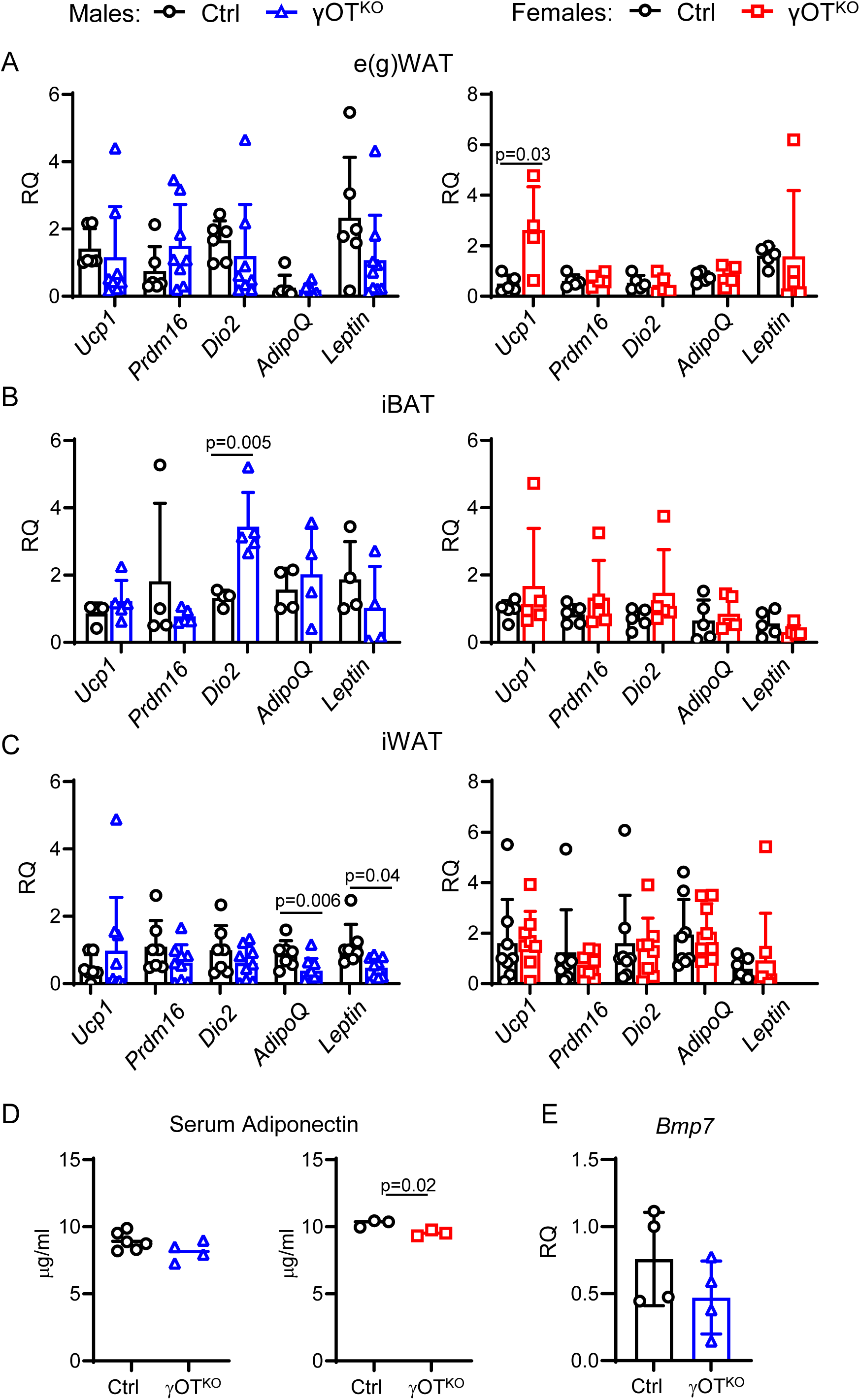
Analysis of metabolic gene markers and adiponectin protein levels in 7 mo old male and female γOT^KO^ and Ctrl mice. A, B, and C. Metabolic gene markers expression in epidydimal/ gonadal WAT (e(g)WAT), interscapular BAT (iBAT), and inguinal WAT (iWAT), respectively. D. Levels of adiponectin in circulation in males and females. E. Expression of *Bmp7* transcript in bone homogenates. Statistical significance of differences between two groups were calculated using unpaired two-tailed student’s t test. Exact and significant p values (< 0.05) are included in the graphs.

However, the levels of circulating adiponectin were decreased in γOT^KO^ females and had a tendency to decrease in γOT^KO^ males (Fig. 2D), which corresponded to a tendency for decreased fat mass shown in Fig. 2D and 2E. Of note, we did not observe changes in *Bmp7* expression in bone of γOT^KO^ animals, which correlated with adipose tissue beiging, as reported previously (Fig. 2E) [10].

### 3.3 γOT^KO^ mice are glucose tolerant but insulin resistant

In both sexes, fasting glucose levels and glucose disposal measured in GTT test were not different from Ctrl. However, blood glucose levels in γOT^KO^ mice measured at the first time point after IP glucose injection (15 min) were either significantly (males) or had a tendency (females) to be lower as compared to Ctrl mice (Fig. 3A). This point of measurement corresponds to an innate cellular disposal of glucose before cells are sensitized to glucose by insulin following its release from pancreas in response to high glucose levels in circulation. Nevertheless, calculated values of the GTT Area Under Curve (AUC) were not different from Ctrl in both sexes (Fig. 3A).

**Figure 3.**
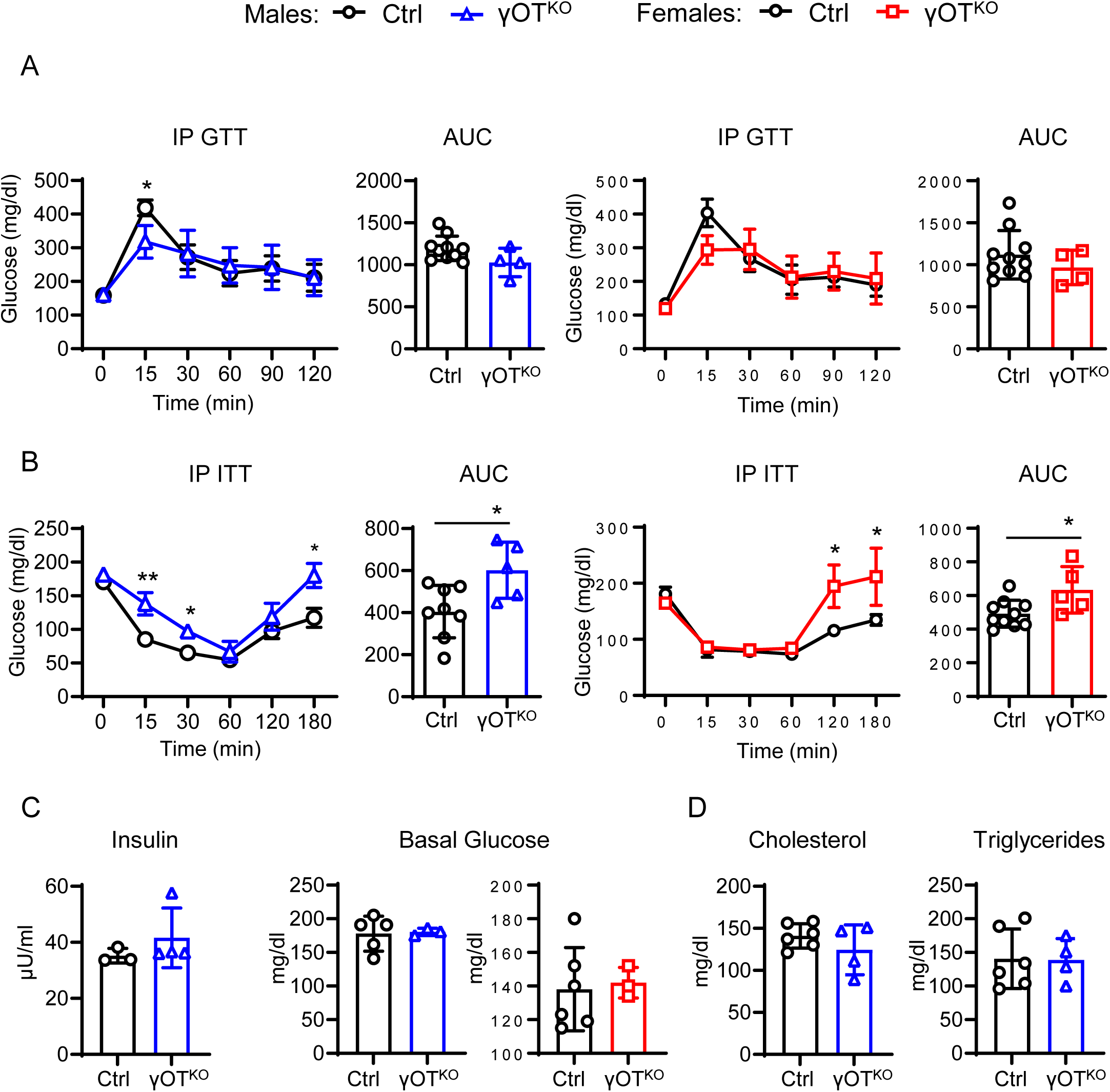
Glucose and lipid metabolism profile of 6 mo old male and female Ctrl and γOT^KO^ mice. A. and B. Intraperitoneal (IP) Glucose Tolerance Test (GTT) and Insulin Tolerance Test (ITT), respectively, for males and females after 4hr fasting. Males – Ctrl: n=11, γOT^KO^: n=5; females – Ctrl: n=11, γOT^KO^: n=5. C. Serum insulin and basal (non-fasted) glucose levels. Ctrl (n=3-5), γOT^KO^ (n=4). D. Serum levels of total cholesterol and triglycerides in males. Ctrl (n=6) and γOT^KO^ (n=4). Statistical significance of differences between two groups were calculated using unpaired two-tailed student’s t test. *p < 0.05; **p < 0.01.

Surprisingly, males, and to a lower degree females, were characterized with apparent insulin intolerance. Glucose load was cleared poorly in γOT^KO^ males, and at the later time points in γOT^KO^ females (Fig. 3B). The calculated values of AUC in ITT assay were significantly higher for γOT^KO^ males and females, as compared to Ctrl (Fig. 3B). Importantly, the apparent insulin intolerance did not correlate with changes in non-fasted serum levels of insulin and glucose in γOT^KO^ males and females, as compared to Ctrl (Fig. 3C). Similarly, lipid profile of γOT^KO^ mice was not affected as serum levels of triglycerides, and cholesterol (total and HDL), as well as liver weight (not shown), were not different from Ctrl mice (Fig. 3D).

These results, together with a lack of changes in extramedullary adipocyte tissue metabolism, suggest that osteocyte metabolism under control of PPARG directly contributes to the systemic energy metabolism, including insulin sensitivity and energy expenditure.

### 3.4 Transcriptomic analysis of *in vivo* osteocytes points to mitochondrial dysfunction in the absence of PPARG

To investigate PPARG role in regulation of osteocyte metabolism and function, we performed high throughput microarray analysis of PPARG-controlled transcriptome using osteocytes freshly liberated from cortical femora bone of 4 mo old γOT^KO^ and Ctrl males (referred later to *in vivo* osteocytes). With a set up threshold for differentially expressed transcripts of two-fold and higher and a False Discovery Rate (FDR) at p < 0.05, this analysis showed that PPARG deficiency leads to upregulation of up to 4,889 and downregulation of 1,876 transcripts (Fig. 4A). To assure confidence in these results, we performed transcriptome analysis of MLO-Y4 clones edited for *Pparγ* with CRISPR/Cas9 (γY4^KO^) and mock-edited control (γY4^Ctrl^), and identified 1,878 transcripts upregulated and 108 transcripts downregulated in γY4^KO^ cells, as compared to γY4^Ctrl^ cells.

**Figure 4.**
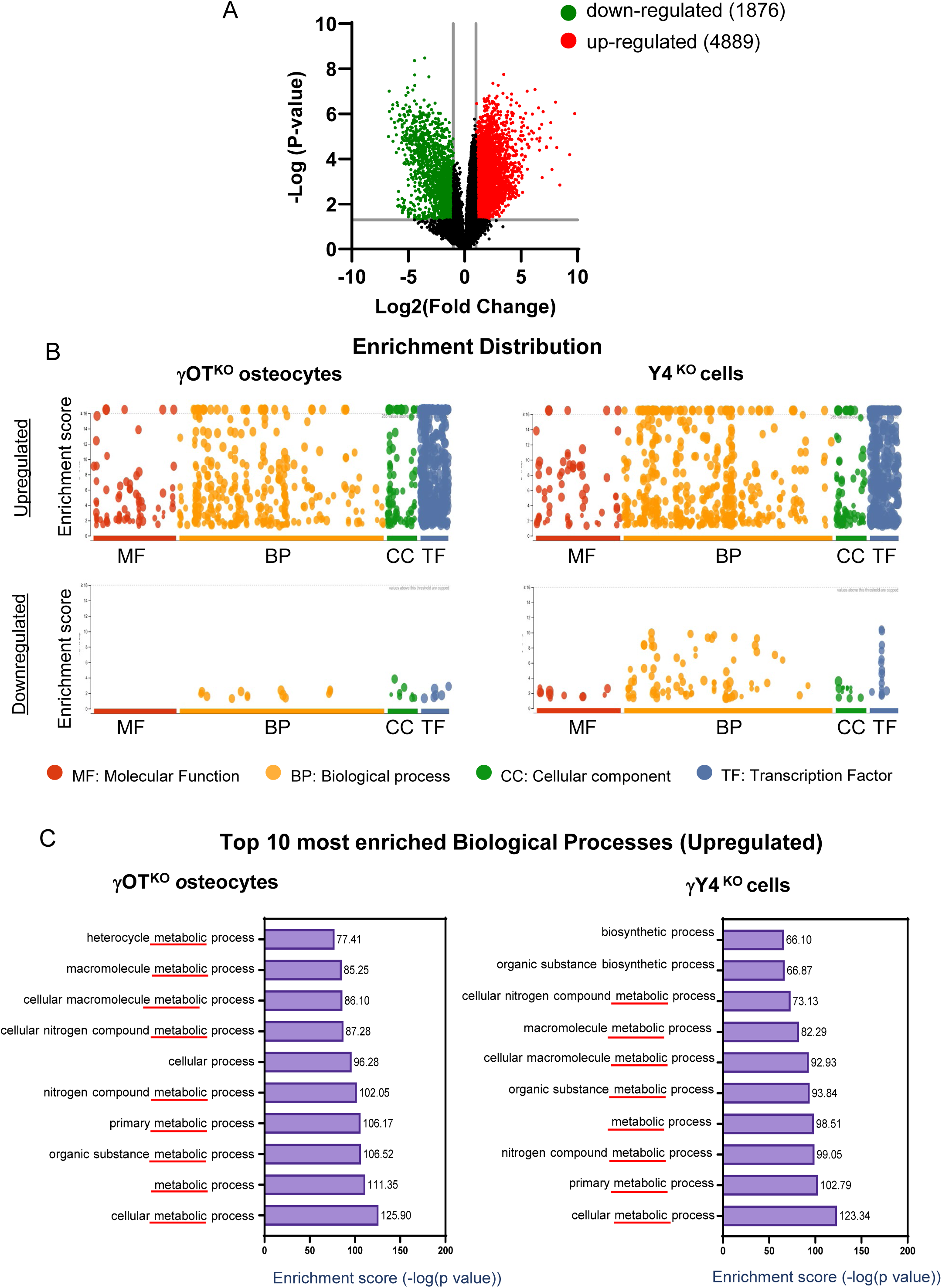
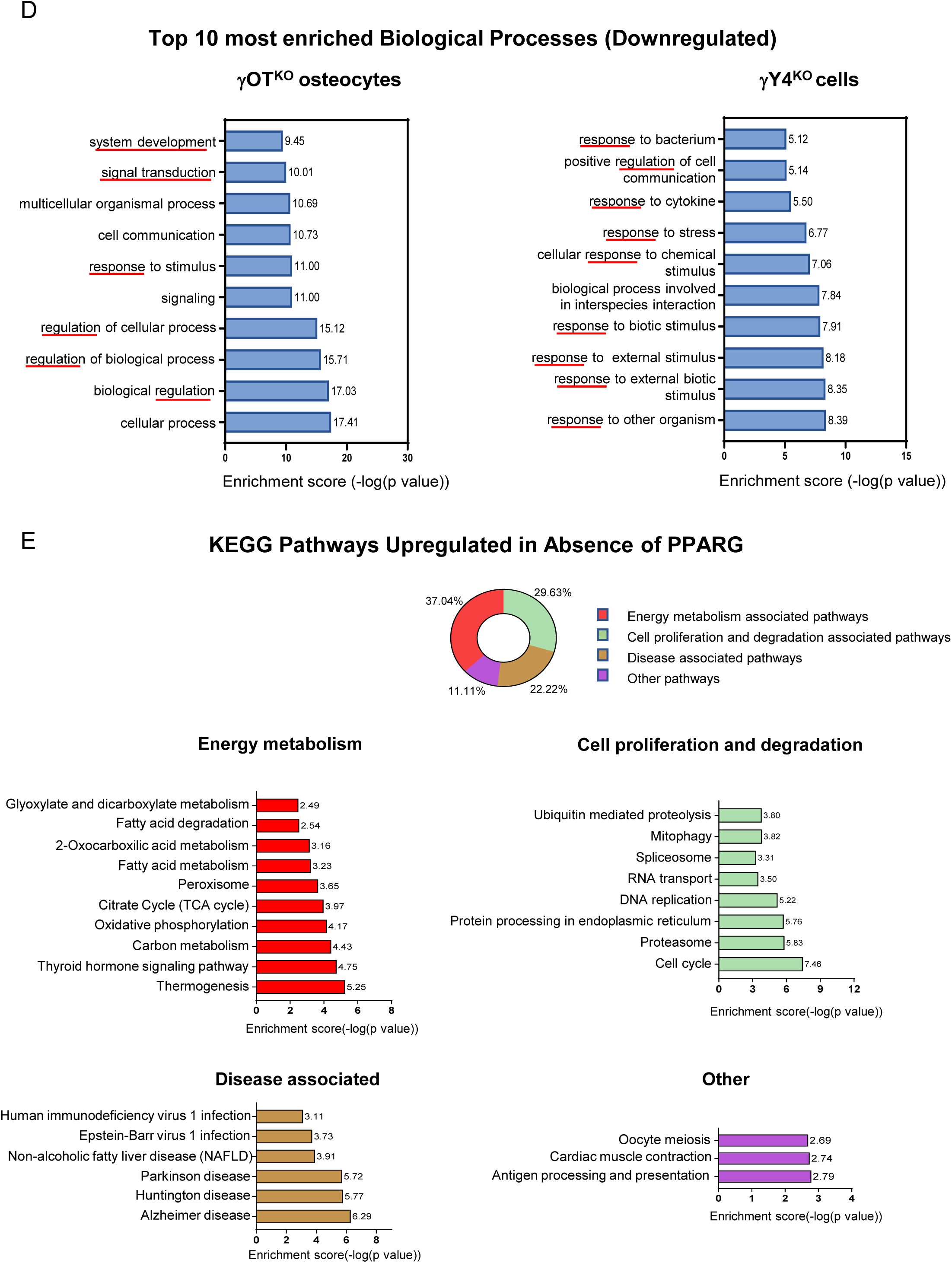
Transcriptome analysis of osteocytes as a function of PPARG. A. Volcano plot of differentially expressed transcript in *in vivo* osteocytes freshly isolated from femora cortical bone of 4 mo old γOT^KO^ and Ctrl male mice. The analysis was performed using the ArrayStar microarray platform as described in Material and Methods. B. GO term enrichment analysis of differentially expressed (fold change>2) statistically significant (p<0.05) genes in freshly isolated osteocytes (γOT^KO^ osteocytes) and MLO-Y4 cells edited for *Pparγ* with CRISPR/Cas9 (γY4^KO^ cells). Graphs were generated using g:Profiler enrichment scores as -log10 (p-value) of upregulated and downregulated genes. C. and D. Top 10 most enriched upregulated and downregulated BP (biological process) GO terms, respectively. E. KEGG analysis of upregulated pathways in γOT^KO^ osteocytes including pie chart of the identified four major pathways categories and -log10 (p-value) enrichment scores of specific pathways in each category.

An analysis of functional clusters using Gene Ontology (GO) enrichment platform and g profiler, was restricted to following categories: molecular function (MF), biological processes (BP), cellular components (CC), and transcription factors (TF). As shown in Fig. 4B, there is a profound representation of upregulated clusters in MF, BP and CC categories in *in vivo* osteocytes and MLO-Y4 cells deficient in PPARG. Similarly, TF category which consists of separate transcripts for transcription factors showed their robust upregulation in *in vivo* osteocytes and γY4^KO^ cells. These analyses together suggest that PPARG in osteocytes acts as an efficient transcriptional repressor for a number of genes, while its function as transcriptional activator of some genes including sclerostin is much less frequent.

The analyses of Biological Processes category showed remarkable similarity between functional clusters of *in vivo* osteocytes and γY4^KO^ cells. The 10 top upregulated clusters in both sets represented cellular metabolic processes with -log(p value) enrichment scores ranging from 70 to 125, which correspond to p values of 10^-70^ and 10^-125^, respectively (Fig. 4C). In contrast, the downregulated top 10 most enriched clusters consisted of regulation of cellular processes, responses to stimulus and signaling with enrichment scores ranging from 5 to 17 which correspond to p values of 10^-5^ and 10^-17^, respectively (Fig. 4D).

An analysis of upregulated pathways representing high-level functions, using Kyoto Encyclopedia of Genes and Genomes (KEGG) [21], had selected 27 pathways that felt into 4 categories: energy metabolism, cell proliferation and degradation, disease associated and other pathways (Fig. 4E). Among them, energy metabolism constituted 37.0% of selected pathways and included clusters for fatty acids metabolism, oxidative phosphorylation, and thermogenesis, with enrichment scores ranging from 2.5 to 5.3 (Fig. 4E). The category of cell proliferation and degradation constituted 29.6% and consisted of pathways associated with mitophagy, nucleic acids and protein metabolism, and cell cycle, with enrichment scores ranging from 3.3 to 7.5 (Fig. 4E).

Overall, the transcriptomic profile of osteocytes deficient in PPARG points to the complex function of this nuclear receptor which can amass to functioning as a “transcriptional molecular break” in cells of osteocytic lineage. The function of this “molecular break” mechanism includes control of osteocyte metabolism and their mitochondrial activity.

### 3.5 PPARG deficiency increases oxidative phosphorylation in γY4^KD^ cells and mitochondrial gene expression in γOTKO osteocytes

High energy metabolism in the absence of peripheral fat depots beiging in γOT^KO^ mice together with osteocyte transcriptomic analysis, suggest a profound effect of PPARG on regulation of osteocyte bioenergetics. Since *in vivo* γOT^KO^ osteocytes indicated robust changes in expression of genes regulating cellular metabolism and an increase in expression of genes involved in mitochondrial ATP production, the mitochondrial activity as a function of PPARG was measured in γY4^KD^ cells using the Agilent Seahorse XF Cell Mito Stress.

A reduction of *Pparγ* transcript by 50% in γY4^KD^ cells (Fig. 5A) led to increased oxidative phosphorylation (OxPhos) measured as oxygen consumption rate (OCR) (Fig. 5B). There was no change in extracellular acidification rate (ECAR) indicating no effect on anaerobic glycolysis and lactate production (Fig. 5C). PPARG reduction resulted in increased basal and maximal cellular respiration and increased ATP production (Fig. 5D). This was associated with a significant increase in spare respiratory capacity and a tendency to increase in proton leak, with no change in non-mitochondrial respiration (e.g. originating from pro-oxidant and pro-inflammatory enzymes) indicating that observed increased respiration is related to mitochondria function (Fig. 5D). Increased binding of tetramethylrhodamine ethyl ester (TMRE) to the polarized mitochondrial membrane in γY4^KD^ cells supported increased mitochondrial activity in γY4^KD^ cells (Fig. 5E).

**Figure 5.**
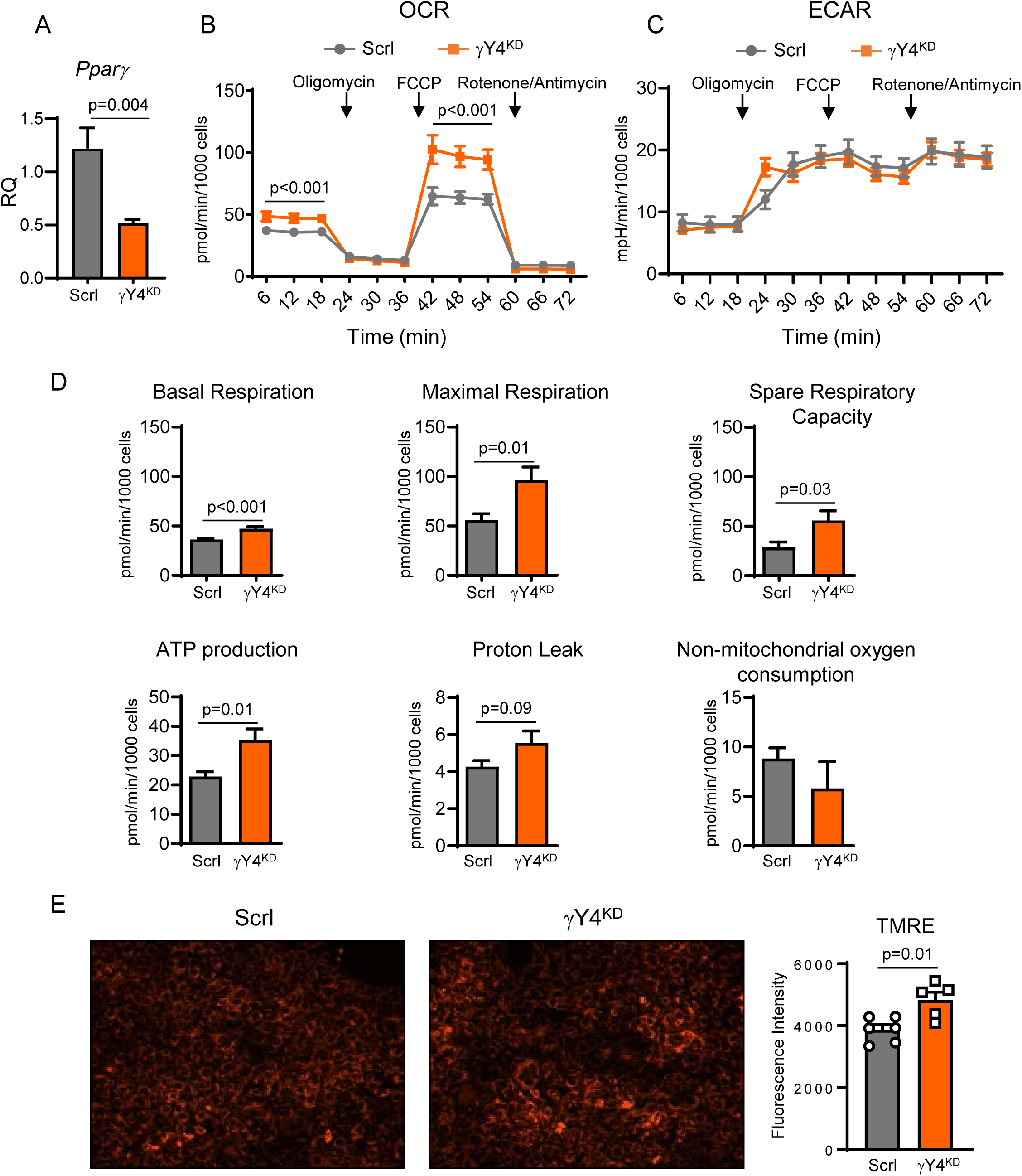
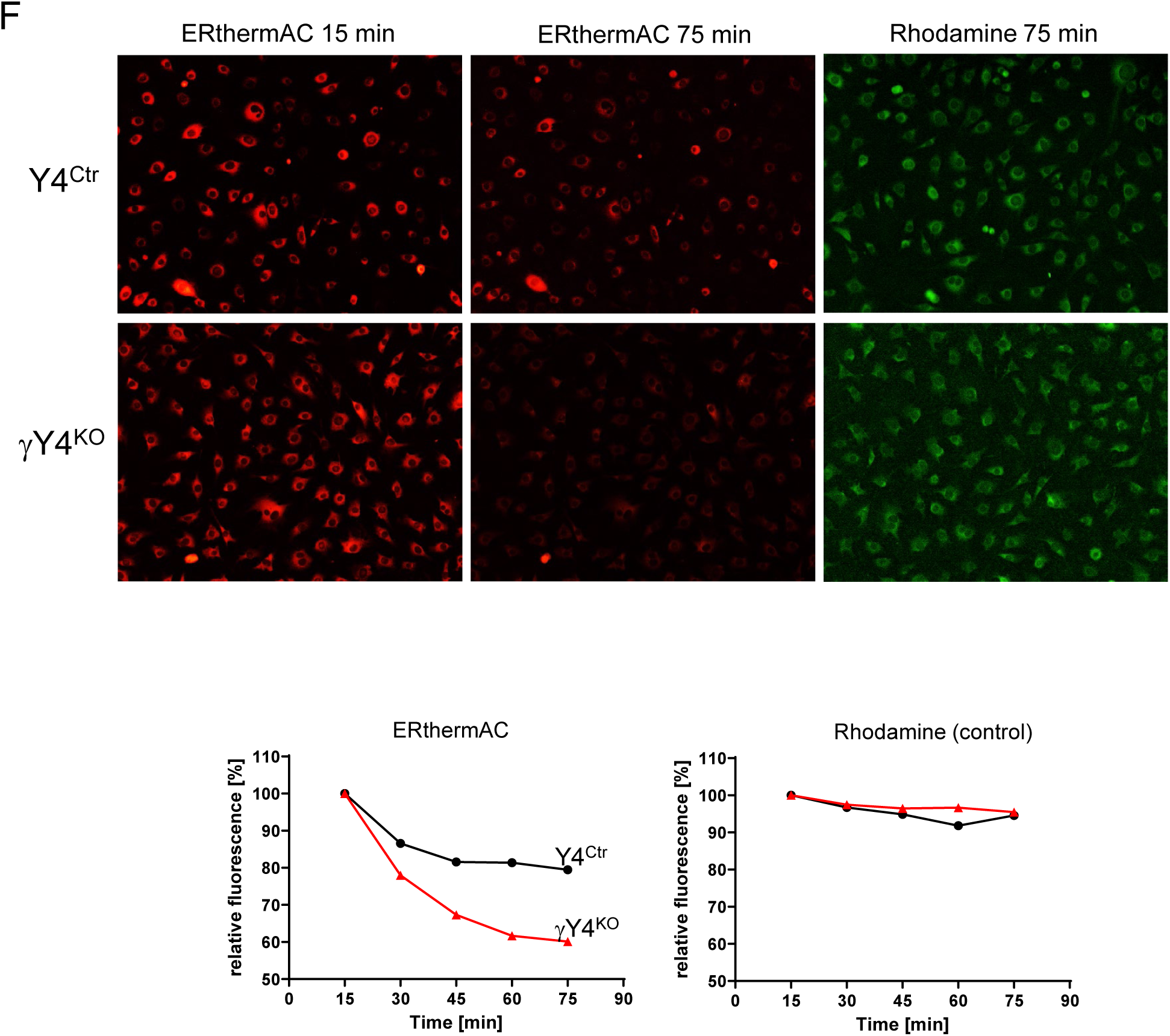
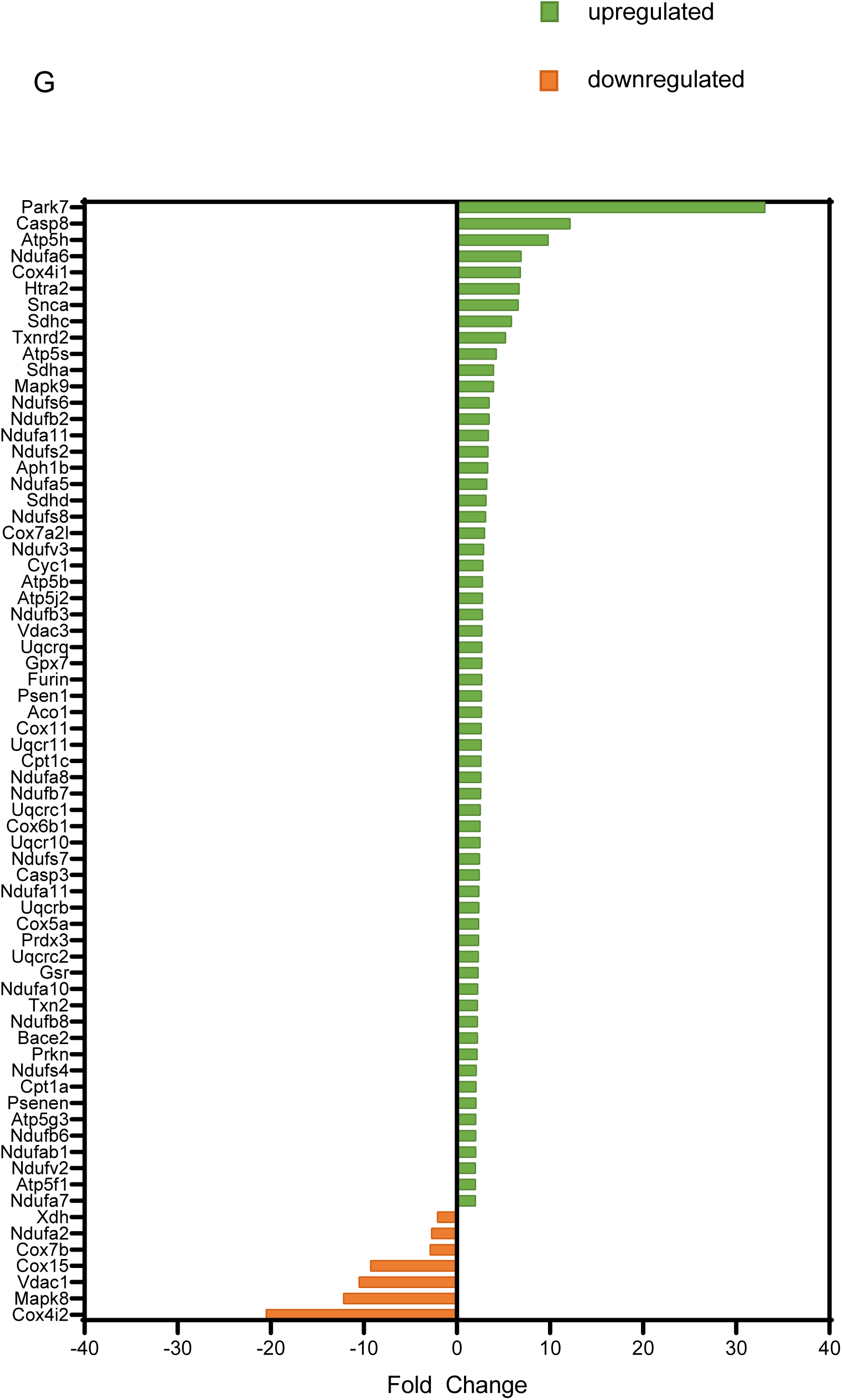
Osteocyte cellular respiration and mitochondrial activity as a function of PPARG. A. Levels of *Pparγ* expression in MLO-Y4 cells after knock-down using siRNA either specific to *Pparγ* (γY4^KD^ cells) or non-specific scrambled control (Scrl cells). (n=3 per group). B. Increased oxygen consumption rate (OCR) in γY4^KO^ cells as compared to Scrl cells. C. Unchanged extracellular acidification rate (ECAR) in γY4^KO^ cells. D. Seahorse MitoStress measurements of mitochondrial function (n=24 well per group). Values were normalized per 1000 cells. E. Mitochondrial membrane potential visualized with TMRE staining (n=5/6 well per group) F. Optical visualization of thermogenesis using ERthermAC fluorescence dye. Stable level of fluorescence observed over the course of the experiment in cells stained with Rhodamine B indicates that loss of fluorescence in cells stained with ERthermAC results from a change of the temperature within the cell, and not resulting from photobleaching or unspecific loss of the dye. Images of cells were acquired with Incucyte S3 Live-Cell Analysis System at 15 minutes intervals using 20X magnification in red channel (567-607 nm excitation, 622-704 emission). G. Differentially expressed genes in γOT^KO^ vs Ctrl *in vivo* osteocytes selected by Ingenuity Pathway Analysis in category of Mitochondrial Dysfunction. Statistical significance of differences between two groups was calculated using unpaired two-tailed student’s t test.

High activity of mitochondria and heat production was confirmed with optical visualization of thermogenesis using ERthermAC, a BODIPY derivative fluorescence dye. ERthermAC forms a contiguous thermometer in the immediate mitochondrial vicinity by targeting the endoplasmic reticulum membrane associated with mitochondria [20]. In this assay, increase in temperature is visualized by a loss of ERthermAC fluorescence as a function of time. As shown in Fig. 5F, mitochondria of γY4^KO^ cells lose the fluorescence at a rate 2-fold faster than γY4^Ctrl^ cells confirming higher mitochondrial heat production in osteocytes deficient in PPARG. Stable level of fluorescence observed over the course of the experiment in cells stained with Rhodamine B indicates that loss of fluorescence in cells stained with ERthermAC results from a change of the temperature within the cell, and not resulting from photobleaching or unspecific loss of the dye.

An observed increase in mitochondrial respiration and ATP production was reflected in a pattern of gene expression regulating electron transport and ATP production (Fig. 5G and Supplementary Table 2). In both, *in vivo* osteocytes of γOT^KO^ mice and γY4^KD^ cells (not shown), there was a significant increase in the expression of enzymes regulating NADH: ubiquinone oxidoreductase activities in Complex I (*Nduf* family), ubiquinone dependent pathway for electrons transport from Complexes I and II to Complex III (*Uqcr* family), Cytochrom C dependent electrons transport from Complex III to IV (*Cox* family), and ATP synthases of Complex IV (*Atp* family). These transcriptional changes amounted to mitochondrial dysfunction in PPARG-deficient osteocytes, as predicted by Ingenuity functional clusters analysis (Supplementary Table 2).

### 3.6 PPARG deficiency increases γY4^KD^ cells usage of glucose, fatty acids and glutamine as fuel for energy production

Changes in mitochondrial activity were associated with changes in fuel use. The PPARG effect on capacity and dependency of MLO-Y4 cells to oxidize three critical mitochondrial fuels: glucose, fatty acids, and glutamine, was measured using the Agilent Seahorse XF Mito Fuel Flex Test. This test determines the rate of oxidation in basal state of each fuel by measuring mitochondrial respiration (OCR) in the presence of selective inhibitors such as UK5099, Etomoxir and BPTES which inhibit respectively glucose, long chain-fatty acids, and glutamine oxidation. The assay measures both fuel capacity which informs of cells’ ability to meet metabolic demand from one fuel when other fuel pathways are inhibited, and fuel dependency which informs of cells’ reliance on a certain fuel to maintain basal respiration. As shown in Fig. 6A, in basal conditions MLO-Y4 cells rely mostly on glucose, and to a lesser extent on fatty acids, as a fuel source. Glutamine use is negligeable. Decreased PPARG activity in γY4^KD^ cells, increases glucose capacity (Fig. 6A), and simultaneously increases fatty acids and glutamine dependency (Fig. 6B).

**Figure 6.**
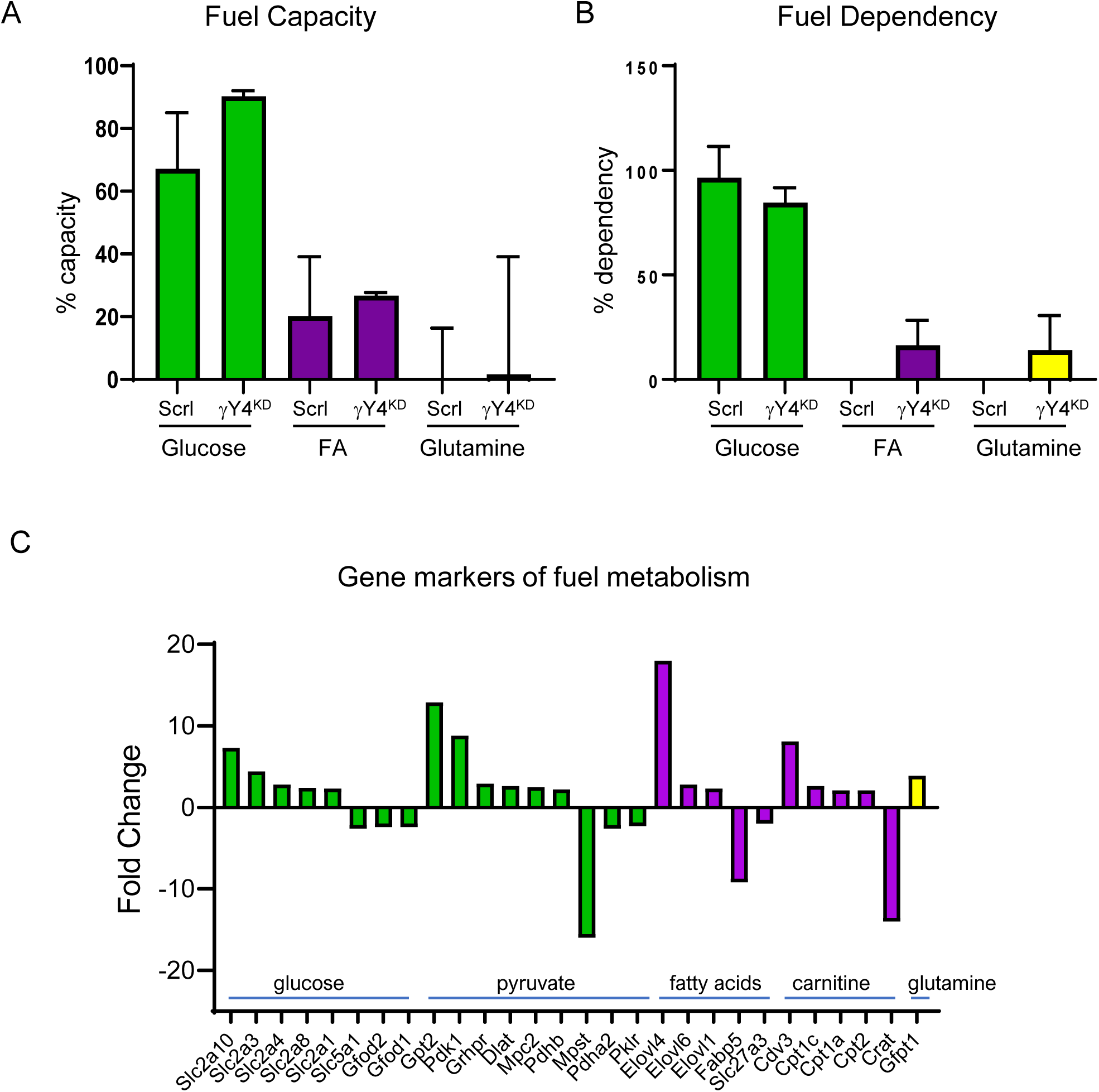
Fuel usage in MLO-Y4 cells edited for *Pparγ* with CRISPR/Cas9 (γY4^KO^) and mock control (Ctrl) cells. A. Fuel capacity measured with Seahorse FuelFlex assay (n=5/6 well per group). B. Fuel dependency measured in the same assay as in A (n=5/6 well per group). C. Fold change in expression of gene markers of fuel metabolism in *in vivo* osteocytes freshly isolated from cortical bone of γOT^KO^ mice as compared to osteocytes isolated from Ctrl mice. Green bars – transcripts related to glucose metabolism; purple bars – transcripts related to fatty acids metabolism; yellow bar – transcript related to glutamine metabolism. Statistical significance in differences between two groups was calculated using unpaired two-tailed student’s t test.

The pattern of changes in fuel use was confirmed in *in vivo* osteocytes deficient in PPARG. As shown in Fig. 6C, *in vivo* γOT^KO^ osteocytes have increased expression of glucose transporters (2.2 - 7.3 folds) including *Slc2a* or *Glut 1, 3, 4, 8,* and *10*, and increased expression of genes coding for proteins essential for pyruvate formation and metabolism (2.2 - 12.9 folds). At the same time, a significant decrease in transcripts involved in alternative metabolic pathways, which include cysteine metabolism (*Mpst*) and glycolysis (*Pklr*), was observed. In respect to fatty acid metabolism, PPARG deficiency increased synthesis of long chain fatty acids (*Elovl 1, 4* and *6*) and enzymes responsible for carnitine formation and its transport to mitochondria (*Cpt1a* and *1c*, and *Cpt2*). Simultaneously, there is a decrease in *Fabp5* responsible for transport of certain fatty acids to nucleus, and fatty acids ligase *Scl27a3*, indicating alterations in fatty acid metabolism. Consistent with increased glutamine dependency, there is a 3.9-fold increase in expression of transcripts for glutamine fructose-6-phosphate transaminase 1 (*Gfpt1*), an enzyme essential for conversion of glutamine to glutamate, which enters TCA cycle.

These findings indicate that PPARG is essential for regulation of osteocyte bioenergetics by acting as a molecular coordinator of glucose, fatty acids and glutamine utilization.

### 3.7 Dysfunctional insulin signaling and decreased insulin sensitivity of PPARG deficient osteocytes

One of PPARG’s essential activities is sensitizing cells to insulin *via* insulin receptor followed by activation of AKT-dependent targets in the insulin signaling pathway. The KEGG analysis of *in vivo* γOT^KO^ osteocyte transcriptome identified a large number of differently expressed transcripts in the functional cluster for insulin signaling pathway (KEGG cluster: mmu04910) (Fig. 7A and 7B). The expression of 83.3% of genes in this cluster was detected in osteocytes, out of which almost 52% was differentially expressed in osteocytes derived from γOT^KO^ mice as compared to Ctrl mice. Among them, transcripts for insulin receptor (*Insr*), insulin receptor substrate (*Irs3*), and PPARG coactivator 1α (*Ppargc1α*) were significantly downregulated indicating decrease in an overall function of this pathway in PPARG deficient osteocytes (Fig. 7A and 7B).

**Figure 7.**
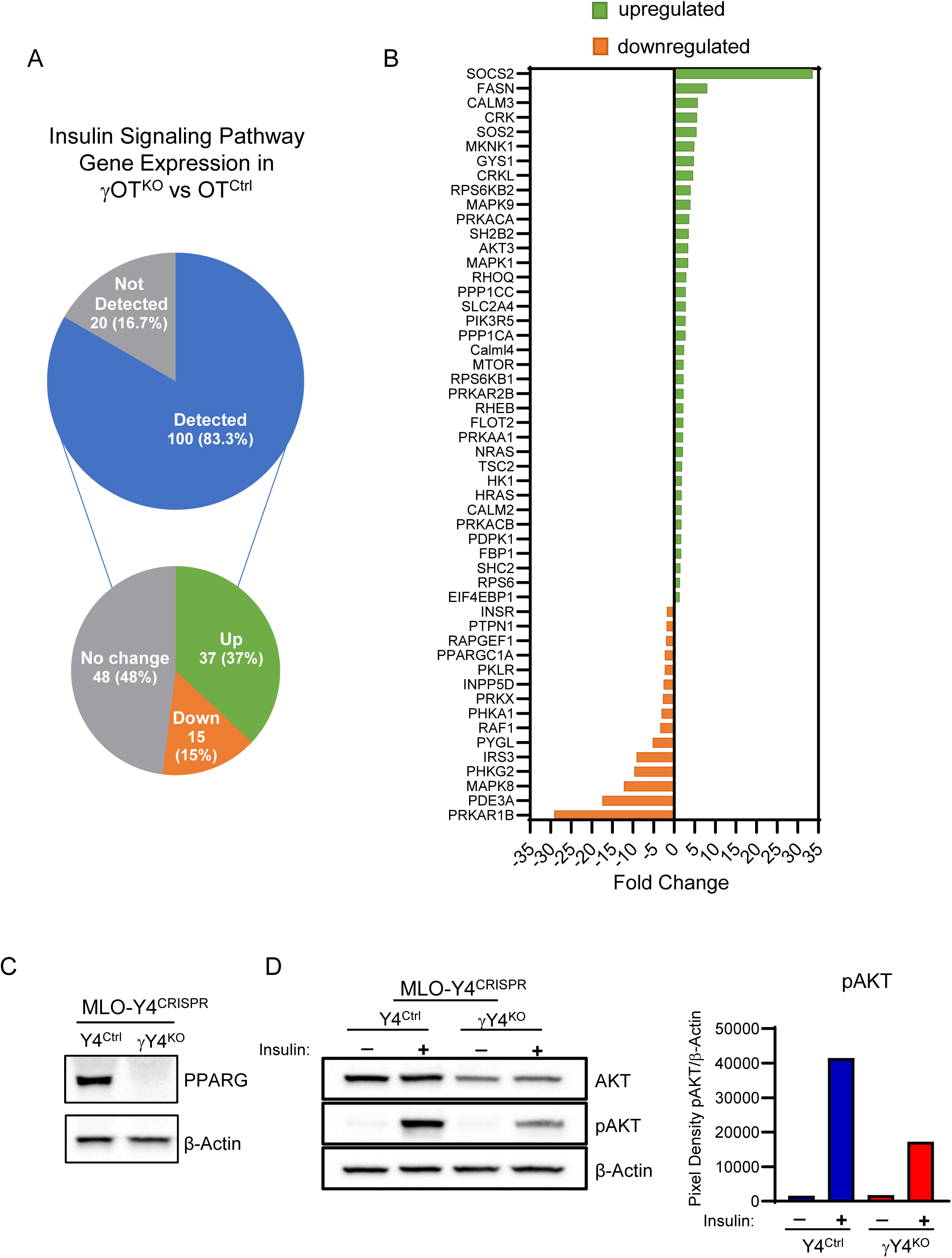
PPARG controls insulin signaling in osteocytes. A. and B. KEGG functional pathway analysis of differentially expressed transcripts in γOT^KO^ vs Ctrl *in vivo* osteocytes. C. Western blot analysis of PPARG protein levels in Y4^Ctrl^ and γY4^KO^ CRISPR/Cas9 edited MLO-Y4 clones. D. Western blot analysis of AKT and Phospho-AKT (pAKT) levels after insulin stimulation of Y4^Ctrl^ and γY4^KO^ clones. Bands density for pAKT were measured with Image J and normalized to β-actin bands density, and plotted in the accompanying graph.

Decreased response to insulin was confirmed in γY4^KO^ cells at the levels of AKT activation. As shown in Fig. 7C and 7D, an absence of PPARG in γY4^KO^ cells decreases overall levels of AKT protein and its phosphorylation (pAKT) in response to insulin stimulation is compromised. These results demonstrate that osteocytes respond to insulin in PPARG dependent fashion and suggest that the systemic insulin resistance seen in γOT^KO^ animals may be at least in part due to insulin resistance in PPARG-deficient osteocytes.

### 3.8 PPARG deficiency in osteocytes increases oxidative stress

Increases in metabolic processes and mitochondrial activity/dysfunction and/or a decrease in insulin signaling (Fig. 7), together with evidence of increased oxidative phosphorylation associated with a tendency to increased proton leak (Fig. 5) and increased fuel flux (Fig. 6), may lead to increased oxidative stress and accumulation of ROS in osteocytes deficient in PPARG. Indeed, two *in vitro* assays confirmed that PPARG deficiency increases oxidative stress in osteocytes. The GSH:GSSG assay which measures SOD activity (Fig. 8A) and DCFDA assay which measures exact ROS production (Fig. 8B), indicated an increase in oxidative stress in γY4^KO^ cells. However, the pattern of gene expression of *in vivo* γOT^KO^ osteocytes showed a significant increase in the transcripts in the Nrf2-mediated Oxidative Stress Ingenuity category (Fig. 8C). The list of upregulated transcripts includes *Sod1* (3.4-fold), *Hmox1* (3.7-fold), *Hmox2* (7.3-fold), and transcripts coding for glutathione transferases, peroxidases, and peroxiredoxins; enzymes essential for cellular defense against ROS. This apparent discrepancy between results from *in vitro* assays indicating decreased SOD activity and accumulation of ROS, and *in vivo* osteocytes from 6 mo old mice showing increased expression of genes in oxidative stress category may be interpreted as a defense response of γOT^KO^ osteocytes which sense an increase in ROS production and are not yet damaged to induce the defense response.

**Figure 8.**
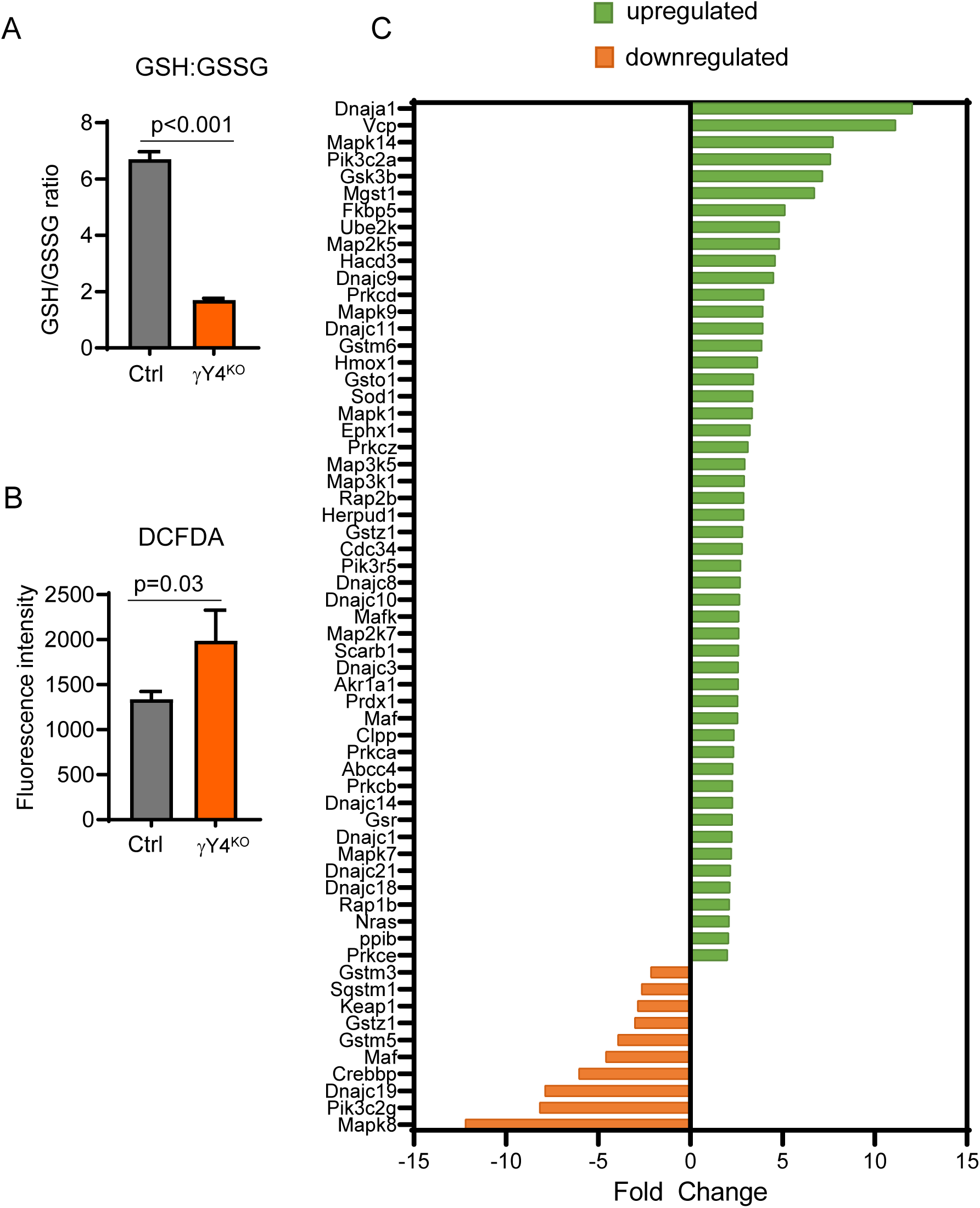
PPARG deficiency in osteocytes increases oxidative stress. A. SOD activity in γY4^KO^ as compared to CRISPR/Cas9 mock-edited cells (C). B. ROS accumulation in γY4^KO^ cells as compared to C cells. C. Expression of transcripts in Nrf2 pathway identified by Ingenuity Pathway Analysis in *in vivo* osteocytes freshly isolated from cortical bone of γOT^KO^ and Ctrl 6 mo old male mice.

## 4. Discussion

In the current study, we provide evidence that energy metabolism and bone physiology are interconnected at the level of osteocytes and that PPARG plays a crucial role in this relationship. Our model of incomplete phenotypic penetrance of osteocyte specific PPARG deficiency inadvertently revealed an important contribution of osteocyte bioenergetics to the levels of systemic energy metabolism. The evidence suggests that PPARG in osteocytes acts as a transcriptional repressor of metabolic activities amounting to the control of mitochondrial activity, ATP production, fuel use, and protection from oxidative stress and ROS accumulation. In addition, these data highlight a likely role of PPARG in control of osteocyte insulin signaling, contributing to the systemic glucose metabolism.

Our model adds a new mechanistic insight to the two other existing models of genetic ablation of PPARG in osteocytes; i.e. the Brun et al. model of PPARG deletion in osteocytes under Dmp1-Cre promoter-driver and the Kim et al. model of PPARG deletion from osteoblast and osteocytes under the Ocn-Cre promoter-driver [9,10]. Both models demonstrated an increase in systemic energy metabolism and increase in insulin sensitivity associated with extramedullary fat beiging, however they differ in proposed possible mechanisms. The Brun et al. model suggested that beiging of extramedullary adipose depots was due to endocrine effects from BMP7 produced in bone. Since the authors did not observe any changes in sclerostin expression, they eliminated this protein from consideration [10]. In contrast, the Kim et al. model suggested that decreased levels of circulating sclerostin have a beiging effect on extramedullary adipose tissue probably by derepressing WNT pathway activity in adipocytes. The authors proved the association of fat beiging with circulating sclerostin levels in a series of loss-of-function and gain-of-function experiments and they confirmed that sclerostin expression is under positive control of PPARG [9]. However, supplementing sclerostin in circulation only partially reverted phenotype indicating that either other circulating factors or other mechanisms, perhaps innate to osteocytes, contributed to this effect.

The γOT^KO^ mice differ in their phenotype from the above models by displaying rather local, not systemic endocrine effects, of PPARG deficiency. This is exemplified by the absence of changes in circulating sclerostin levels. We previously showed that PPARG is a positive transcriptional regulator of sclerostin and its deletion results in a lack of sclerostin transcript and protein in osteocytes [7]. Reduced skeletal sclerostin is supported by the high bone mass phenotype. Both male and female γOT^KO^ mice have increased bone mass, especially in cortical bone which correlates with high osteoblastic activity on the endosteal and periosteal surface, and decreased bone marrow adipose tissue volume for which sclerostin acts as a positive regulator [7,22]. However, as we have shown before and here, sclerostin levels in circulation are not affected, which correlated with a lack of beiging of extramedullary fat. Nevertheless, male γOT^KO^ mice have a high energy phenotype without metabolic changes in extramedullary fat depots. We believe the culprit for this phenotype is increased mitochondrial activity reflected in increased oxidative phosphorylation and ATP production. The increase in mitochondrial activity amounted to their dysfunction due to perturbation in expression of number proteins involved in electron transport despite downregulation of PGC1α coactivator. Our results are consistent with the well documented role of PPARG to control mitochondrial biogenesis and expression of the electron transport chain components either independently or in conjunction with the PGC1α coactivator.

Mitochondrial dysfunction in the absence of PPARG leads to oxidative stress accentuated by increased ROS production and decreased SOD activity in γY4^KO^ osteocytic cells. ROS are known to trigger antioxidant response predominantly by activation of NRF2/KEAP1/ARE pathway leading to the expression of cytoprotective (TXNRD1 and SOD1) and phase II detoxification (HMOX1, FTL1 and GSTP1) enzymes, reliant on reciprocal regulation of PPARG and NRF2 [23]. We have demonstrated that *in vivo* γOT^KO^ osteocytes of relatively young 6 mo old males have significantly increased expression of transcripts in the NRF2/KEAP1/ARE pathway, probably in response to mounting oxidative stress in bone. It remains to be established whether with aging the protective antioxidative response of PPARG-null osteocytes is sustained. Notably, PPARG is known as a powerful defender against oxidative damage induced by ROS [23–26]. Therefore, there is a possibility that with aging, in the absence of PPARG and a lack of control of mitochondrial activity, an oxidative stress may overcome defensive mechanisms and may eventually accelerate osteocyte senescence [27,28] leading to osteocyte dysfunction and weakening of bone material properties [26].

In addition, and in contrast to Brun et al. and Kim et al. models, γOT^KO^ mice have affected glucose disposal in response to intraperitoneally injected insulin without changes in glucose tolerance. This indicates that in γOT^KO^ mice osteocytes do not respond to insulin, the response of which is under control of PPARG [29]. Insulin intolerance of γOT^KO^ mice is in line with insulin intolerance observed in other models of tissue-specific PPARG deletion, such as in muscle or adipose tissue [30,31]. Notably, Brun et al. showed that glucose influx to bone is significantly increased [10]. Indeed, our model showed increased expression of glucose, fatty acids and glutamine transporters and alterations in fuel handling, such as increase in glucose capacity, and increase in fatty acids and glutamine dependency. These point to PPARG being an essential regulator of osteocyte insulin signaling and fuel utilization with global impact. Taken together, we conclude that under control of PPARG osteocyte bioenergetics significantly complement osteocyte endocrine effects in contribution to the systemic energy metabolism.

Interestingly, although bone phenotype of γOT^KO^ female and male mice does not differ, they exhibit sexual divergence in respect to energy metabolism phenotype. Males display high energy phenotype early, while females’ energy metabolism phenotype is not affected except for developing modest insulin resistance later in life. Several possibilities can be discussed, all of them speculative at this point. One is based on a known competition between PPARG and estrogen receptor (ER) for common modulatory proteins including SRC and p300 coactivators. Thus, an absence of PPARG may strengthen activity of ER pathway due to increased availability of common coactivators. Another possibility is that in our model the penetration of the energy metabolism phenotype is much lower in females than in males. Of note, in two other published models, exclusively male animals were employed to illustrate the elevated energy metabolism [9,10]. Thus, it is possible that antagonism between PPARG and ER plays an important role in modulating the osteocyte bioenergetics, in a similar way as in adipocytes which respond to estrogen deficiency by PPARG activation and adipose tissue expansion [32]. It would be of interest to test whether energy metabolism and bioenergetics of osteocytes are increased in estrogen deficient γOT^KO^ females.

Our study of the γOT^KO^ model has its strengths and limitations. The strength consists of thorough analysis of both sexes which revealed divergence in male and female osteocyte phenotype with PPARG deficiency modulated in vivo by sex-specific humoral and innate factors. On the other hand, our model is limited in the absence of PPARG-controlled osteocyte endocrine effects which we believe is a result of limited penetration of PPARG deletion phenotype. It is commonly acknowledged that Cre-loxP cell-specific recombination system is posing uncontrolled complications, which are associated with innate capabilities of the cell to decrease activity or silence foreign transcriptionally active components [18]. However, this limitation unveiled a new role of PPARG in regulation of osteocyte bioenergetics and its contribution to the levels of systemic energy metabolism. These findings led us to the conclusion that PPARG in osteocytes acts as molecular break for regulation of mitochondria activity and its absence dysregulates mitochondria leading to eventual metabolic dysfunction.

In summary, our evidence strongly supports the role of PPARG in osteocytes as a regulator of bone and systemic energy metabolism by controlling osteocyte bioenergetics, in addition to its endocrine activity. Our model of PPARG deletion in osteocytes underscores the intrinsic role of PPARG activity as a molecular break for osteocyte bioenergetics. This study showed that osteocyte energy production is of such magnitude, due to both the sheer number of osteocytes and their high rate of bioenergetics, that it significantly contributes to the overall energy metabolism levels. Our model, together with other models, provides a comprehensive insight for the skeleton acting as a body “energostat” via PPARG activity in osteocytes. These findings are of potential therapeutic interest to develop means of treating bone and metabolic diseases simultaneously by targeting PPARG in osteocytes.

## Authors Contribution

SB, BLC – conceptualization, data curation, writing original draft; SB - investigation; SB, PJC, EC, JL - formal analysis, methodology, validation, visualization; BLC, SB, PJC, MPK, JL, CJR, PRG – writing, reviewing, editing.

## Acknowledgments

This study was supported to BLC and PRG by the National Institute on Aging grant number R01AG071332.

## Abbreviations

PPARG: Peroxisome Proliferator-Activated Receptor Gamma protein
*Pparγ*: either murine gene or transcript coding for PPARG protein
CLAMS: Columbus Laboratory Animal Metabolic System
Ip ITT: Intraperitoneal Insulin Tolerance Test Ip
GTT: Intraperitoneal Glucose Tolerance Test
ROS: Reactive Oxygen Species

