## Supplemental Table 1 for "PPARG in osteocytes controls cell bioenergetics and systemic energy metabolism independently of sclerostin levels in circulation"

Supplementary Table 1. Primers used for real time PCR

| Transcript | Forward | Reverse |
| --- | --- | --- |
| *Ucp1* | GGA TGG TGA ACC CGA CAA CT | AAC TCC GGC TGA GAA GAT CTT G |
| *Prdm16* | CCT AAC TTT CCC CAC TCC CTC TA | GCT CAG CCT TGA CCA GCA A |
| *Dio2* | AAA TGA CCC CTT TGG TTT CC | TTC CCC ATT ATC CCT TTT CC |
| *Sost* | CCT CCT CCT GAG AAC AAC CA | ACA TCT TTG GCG TCA TAG GG |
| *Adipoq* | GGC CGT TCT CTT CAC CTA CG | TGG AGG AGC ACA GAG CCA G |
| *Lep* | ATT TCA CAC ACG CAG TCG GTA T | GGT GAA GCC CAG GAA TGA AG |
| *Bmp7* | ACG GAC AGG GCT TCT CCT AC | ATG GTG GTA TCG AGG GTG GAA |
