## Supplemental Table 2 for "PPARG in osteocytes controls cell bioenergetics and systemic energy metabolism independently of sclerostin levels in circulation"

Supplemental Table 2. Ingenuity Pathway Analysis of differentially expressed genes in *in vivo* γOT^KO^ osteocytes in canonical pathways for Mitochondrial Dysfunction and Oxidative Phosphorylation

| Ingenuity Canonical Pathway | -log(p-value)  (1.3 equals p=0.05;  3 equals p=0.001) | Ratio | Molecules |
| --- | --- | --- | --- |
| Mitochondrial Dysfunction | 6.51 | 0.409 | ACO1, APH1B, ATP5F1B, ATP5MC3, ATP5MF, ATP5PB, ATP5PD, BACE2, CASP3, CASP8, COX11, COX15, COX4I1, COX4I2, COX5A, COX6B1, COX7A2L, COX7B, CPT1A, CPT1C, CYC1, DMAC2L, FURIN, GPX7, GSR, HTRA2, MAPK8, MAPK9, NDUFA1, NDUFA10, NDUFA11, NDUFA2, NDUFA5, NDUFA6, NDUFA7, NDUFA8, NDUFAB1, NDUFB2, NDUFB3, NDUFB6, NDUFB7, NDUFB8, NDUFS2, NDUFS4, NDUFS6, NDUFS7, NDUFS8, NDUFV2, NDUFV3, OGDH, PARK7, PRDX3, PRKN, PSEN1, PSENEN, SDHA, SDHC, SDHD, SNCA, TXN2, TXNRD2, UQCR10, UQCR11, UQCRB, UQCRC1, UQCRC2, UQCRQ, VDAC1, VDAC3, XDH |
| Oxidative Phosphorylation | 4.54 | 0.413 | ATP5F1B, ATP5MC3, ATP5MF, ATP5PB, ATP5PD, COX11, COX15, COX4I1, COX4I2, COX5A, COX6B1, COX7A2L, COX7B, CYC1, DMAC2L, NDUFA1, NDUFA10, NDUFA11, NDUFA2, NDUFA5, NDUFA6, NDUFA7, NDUFA8, NDUFAB1, NDUFB2, NDUFB3, NDUFB6, NDUFB7, NDUFB8, NDUFS2, NDUFS4, NDUFS6, NDUFS7, NDUFS8, NDUFV2, NDUFV3, SDHA, SDHC, SDHD, UQCR10, UQCR11, UQCRB, UQCRC1, UQCRC2, UQCRQ |
